# Tissue-specific multiOMICs analysis of atrial fibrillation

**DOI:** 10.1101/2020.04.06.021527

**Authors:** Ines Assum, Julia Krause, Markus O. Scheinhardt, Christian Müller, Elke Hammer, Christin S. Börschel, Uwe Völker, Lenard Conradi, Bastiaan Geelhoed, Tanja Zeller, Renate B. Schnabel, Matthias Heinig

**Author notes:** I.A. and J.K. contributed equally to this work. T.Z., R.S. and M.H. contributed equally to this work. Correspondence to: M.H., R.B.S.

## Abstract

Genome-wide association studies (GWAS) for atrial fibrillation (AF) have uncovered numerous disease-associated variants. Their underlying molecular mechanisms, especially consequences for mRNA and protein expression remain largely elusive. Thus, novel multiOMICs approaches are needed for deciphering the underlying molecular networks. Here, we integrated genomics, transcriptomics, and proteomics of human atrial tissue which allowed for identifying widespread effects of genetic variants on both transcript (cis eQTL) and protein (cis pQTL) abundance. We further established a novel targeted trans QTL approach based on polygenic risk scores to identify candidates for AF core genes. Using this approach, we identified two trans eQTLs and four trans pQTLs for AF GWAS hits, and elucidated the role of the transcription factor NKX2-5 as a link between the GWAS SNP rs9481842 and AF. Altogether, we present an integrative multiOMICs method to uncover trans-acting networks in small datasets and provide a rich resource of atrial tissue-specific regulatory variants for transcript and protein levels for cardiovascular disease gene prioritization.

## INTRODUCTION

Genome-wide association studies (GWAS) have discovered thousands of disease-associated single nucleotide polymorphisms (SNPs) and improved our understanding of the genetic and phenotypic relationships.^[1]^ In this regard, GWAS have been applied to investigate atrial fibrillation (AF), which affects more than 30 million individuals worldwide.^[2]^ Various genetic risk loci for AF have been identified^[3, 4, 5]^ and have been integrated into genome-wide polygenic risk scores (PRS) for AF risk prediction.^[6, 7]^ However, AF risk loci known so far explain less than half of the estimated disease heritability.^[3]^ In addition, more than 95% of these GWAS variants are localized in noncoding regions^[3]^ not directly affecting the protein sequence but possibly acting through gene-regulatory mechanisms which are mostly unknown.

A common approach to identify target genes of GWAS variants is to consider tissue-specific cis-acting expression quantitative trait loci (eQTL), where genetic variants affect the transcription of nearby genes. However, cis eQTLs only explain a fraction of the identified AF risk loci. Therefore, trans eQTLs, where the variant is distant to the target gene, and more complex genetic or epigenetic mechanisms need to be considered.^[6, 5, 8, 9]^ To date, it remains diffcult to quantify the contribution of cis and trans variants to the heritability of complex diseases such as AF.

Recently, the contribution of trans effects to the genetic architecture of complex traits was theoretically assessed by the omnigenic model.^[10]^ Based on this model, it was estimated that trans genetic effects explain at least 70% of the disease heritability by indirect propagation through gene regulatory networks.^[10]^ Within these networks, multiple trans effects can accumulate on just a few central genes, so-called core genes, which in turn are functionally related to a phenotype. Identifying those core genes by trans eQTLs remains challenging due to the small effect size of each individual locus^[11, 12]^ and the associated large multiple testing burden. Since a PRS summarizes the genetic risk information, it can act as a proxy for the accumulation of trans effects in one score.^[12]^ By correlating the score with transcript expression (eQTS), the propagation of trans effects to mRNA level^[12]^ can be evaluated. However, not only transcript abundance, but also the abundance of translated proteins can determine phenotypic consequences. To date, little is known about genetic effects on protein levels (pQTLs), e.g. through post-transcriptional regulation,^[10, 13, 14, 15, 16]^ especially in atrial tissues. It is both challenging and important to establish methods to identify AF core genes and to integrate data from multiple OMICs levels to improve the understanding of genotype-phenotype relationships.

Here, we present a multiOMICs analysis that uses genomics, transcriptomics, and proteomics of human atrial tissue to better understand how genetics are related to molecular mechanisms of AF. The first aim was to systematically integrate OMICs data and identify genome-wide cis-regulatory mechanisms on transcript as well as protein level. We reasoned that core genes are key for a better understanding of complex molecular pathomechanisms of AF. Therefore, the second aim was to identify candidate core genes for AF and the trans-acting regulatory networks that link them to AF GWAS loci. We developed a novel approach combining the correlation of gene expression with a PRS for AF^[7, 12]^ and pathway enrichment analysis to identify AF-associated biological processes. Based on those processes, candidate core genes for targeted trans QTL analyses with AF GWAS SNPs were selected. This approach allowed the identification of putative core genes, their molecular networks and downstream consequences in AF.

## RESULTS

### Cis QTL analysis

We analyzed disease-independent effects of genetic variants on transcript (cis eQTLs) and protein levels (cis pQTL) of nearby genes. All cis QTLs were calculated using expression values for 16 306 genes and 1 337 proteins (Table 1) with additional PEER factors^[17, 18]^ to adjust for relevant covariates such as technical confounders and cardiovascular risk factors (see methods and Suppl Figure S1-S2). We assessed the replication rate of our eQTLs in GTEx^[19]^ atrial appendage tissue. Effect sizes for the best eQTL (P<1 × 10^−5^) per gene showed a correlation of (P=3.6 × 10^67^) in GTEx, 66% replicated (GTEx P<1 × 10^−5^) and 88% showed concordant allelic effects (see also Suppl “Cis QTL replication”, Suppl Figure S3). Furthermore, correlations between transcript and protein abundances were comparable to previous studies^[14]^ (Suppl Figure S4, Suppl Table S1) indicating high quality of the proteomics measurements.

**Table 1:**
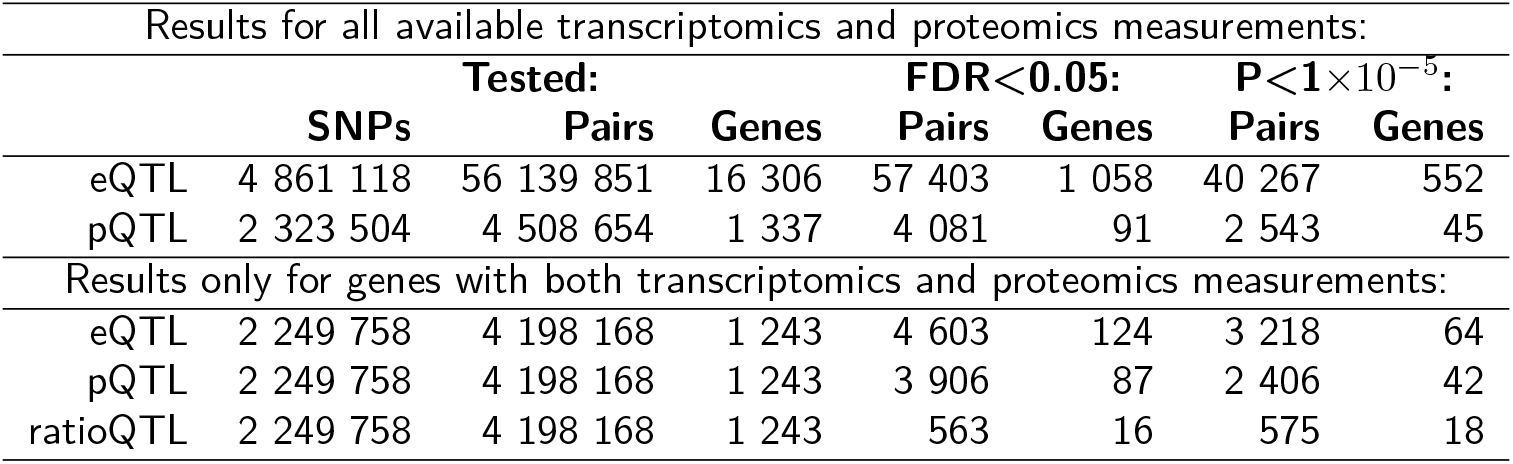
Summary of tested data and discovered QTLs. Results for a FDR<0.05 (according to Benjamini-Hochberg procedure) and P value <1 × 10^−5^. FDR, false discovery rate; eQTL, expression quantitative trait loci; pQTL, protein quantitative trait loci; ratioQTL, ratio quantitative trait loci;

### Cis-regulatory patterns specific to atrial tissue

We first sought to functionally characterize the cis-regulatory variants. Local genetic variation can lead to different modulations in mRNA and protein abundance which are commonly attributed to transcriptional and post-transcriptional regulation.^[14, 15, 20]^ Protein abundances are suggested as more direct determinants for phenotypic consequences of expression QTLs^[14]^ emphasizing the need to integrate mRNA and proteomic measurements to better understand functional genotype-phenotype relationships. Thus, only for the following characterization of cis genetic variation on transcript and protein, we focused on genes with both transcriptomics and proteomics measurements (1 243 genes, Table 1). As observed previously,^[14, 15, 20]^ significant eQTLs and pQTLs (FDR<0.05) differ considerably, as only 8.2% of significant SNP-gene associations are shared between mRNA and proteins (Figure 1, Suppl Figure S5, Suppl Table S2).

**Figure 1:**
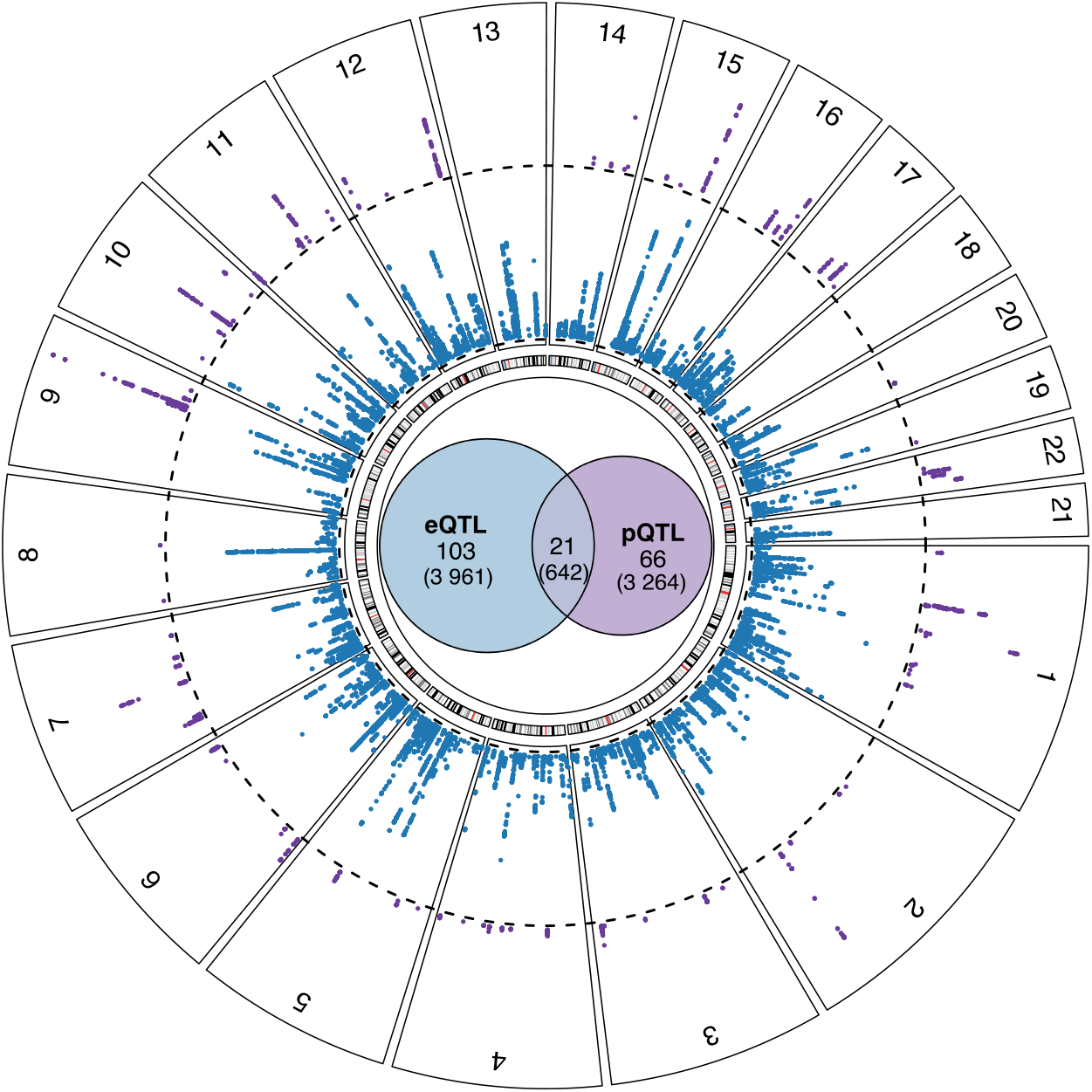
Significant cis eQTLs, cis pQTLs and their overlap. Depicted are the significant cis eQTLs (blue) and pQTLs (purple) at a FDR cutoff of 0.05 (dotted line). Considering only genes with both transcriptomics and proteomics measurements, we visualized the overlap of significant eQTLs and pQTLs in the circle center. In total, 124 and 87 genes had at least one significant eQTL or pQTL, respectively. Only 21 of those genes had at least one overlapping eQTL and pQTL. The numbers in brackets represent the number of significant SNP-gene pairs. eQTL, expression quantitative trait loci; pQTL, protein quantitative trait loci; FDR, false discovery rate;

Divergence of mRNA from protein abundance can arise through diverse molecular mechanisms which we additionally analyzed by calculating protein-per-mRNA ratios (ratioQTLs) (see methods). To identify shared as well as independent effects on mRNA and protein abundance, three simple regulatory categories were analyzed (see methods, Figure 2, Suppl Figure S6, Suppl Table S3):

i. For 11 genes, 430 shared eQTLs / pQTLs were identified, where SNPs affect both mRNA expression and the respective protein abundance as depicted in Figure 2a. The corresponding variants were primarily enriched in cis-regulatory elements such as active transcription start sites (TssA) (Suppl Figure S6a, Suppl Figure S7).
ii. For 37 genes, 1 593 independent eQTLs were identified, where only transcript levels are associated, but the respective protein abundance is independent of the genotype (Figure 2b). Corresponding variants were enriched in elements regulating transcription, e.g. transcription factor binding site (TF BS) or enhancer regions, and within splicing regions (Suppl Figure S6b, Suppl Figure S7a). Possible mechanisms involved in unchanged protein levels remain largely elusive and range from adaptation of translational rate to protein degradation and long-noncoding RNAs.^[21, 22]^
iii. For 21 genes, 1 083 independent pQTLs were identified, where the SNP affects only protein abundance (Figure 2c). pQTL variants were enriched for exonic regions and although not significantly, in binding sites of RNA binding proteins (RBP) (Suppl Figure S6c, Suppl Figure S7a), where they may influence mRNA translation resulting in an independent pQTL association.^[23]^

**Figure 2:**
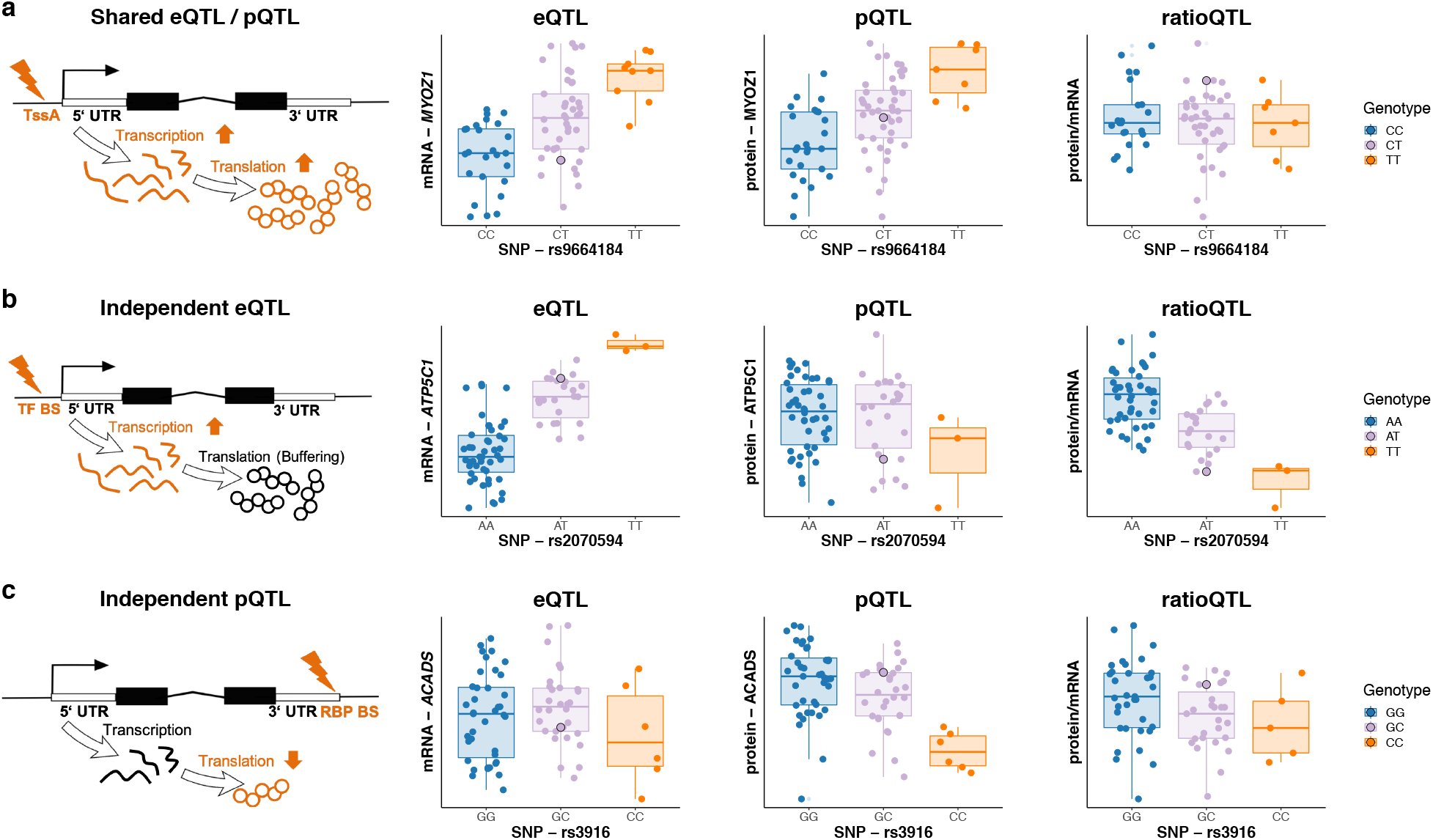
Different genetic regulatory patterns derived by multiOMICs QTL integration. a: Shared eQTLs / pQTLs represent QTLs, where the effect of transcriptional regulation translates into mRNA and protein abundances exemplified by the significant SNP - gene pair rs9664184 - MYOZ1. No corresponding ratio QTL can be observed as the genetic variation is shared across both OMICs levels. b: Independent eQTLs depict variants with regulation on mRNA but not on protein level displayed by the significant SNP - transcript pair rs2070594 - *ATP5C1.* c: Independent pQTLs represent variants that show regulation only on protein level as shown for the SNP-protein pair rs3916 - ACADS. Genetic influence is not observable on transcript level. In the boxplots, the lower and upper hinges correspond to the first and third quartiles (the 25th and 75th percentiles). The median is denoted by the central line in the box. The upper/lower whisker extends from the hinge to the largest/smallest value no further than 1.5 IQR (interquartile range) from the hinge. eQTL, expression quantitative trait loci; pQTL, protein quantitative trait loci; ratioQTL, ratio quantitative trait loci; TssA, active transcription start site; UTR, untranslated region; TFBS, transcription factor binding site; RBP, RNA binding protein;

Altogether, we confirmed that QTL variants corresponding to different categories tended to cluster in distinct genomic regions (Suppl Figure S7, Suppl “Enrichment of functional elements”).^[14, 18, 24]^ By integrating matched transcriptome and proteome data, we were able to differentiate functional regulatory mechanisms not observable by transcriptomics only.

### GWAS overlap and enrichment

In order to investigate genotype-phenotype relationships in the context of cardiovascular disease, we used all available cis QTL data (not only those quantified on both OMICs levels, FDR<0.05) to annotate GWAS variants for phenotypes either related to cardiovascular measurements or cardiovascular disease (Figure 3a).

**Figure 3:**
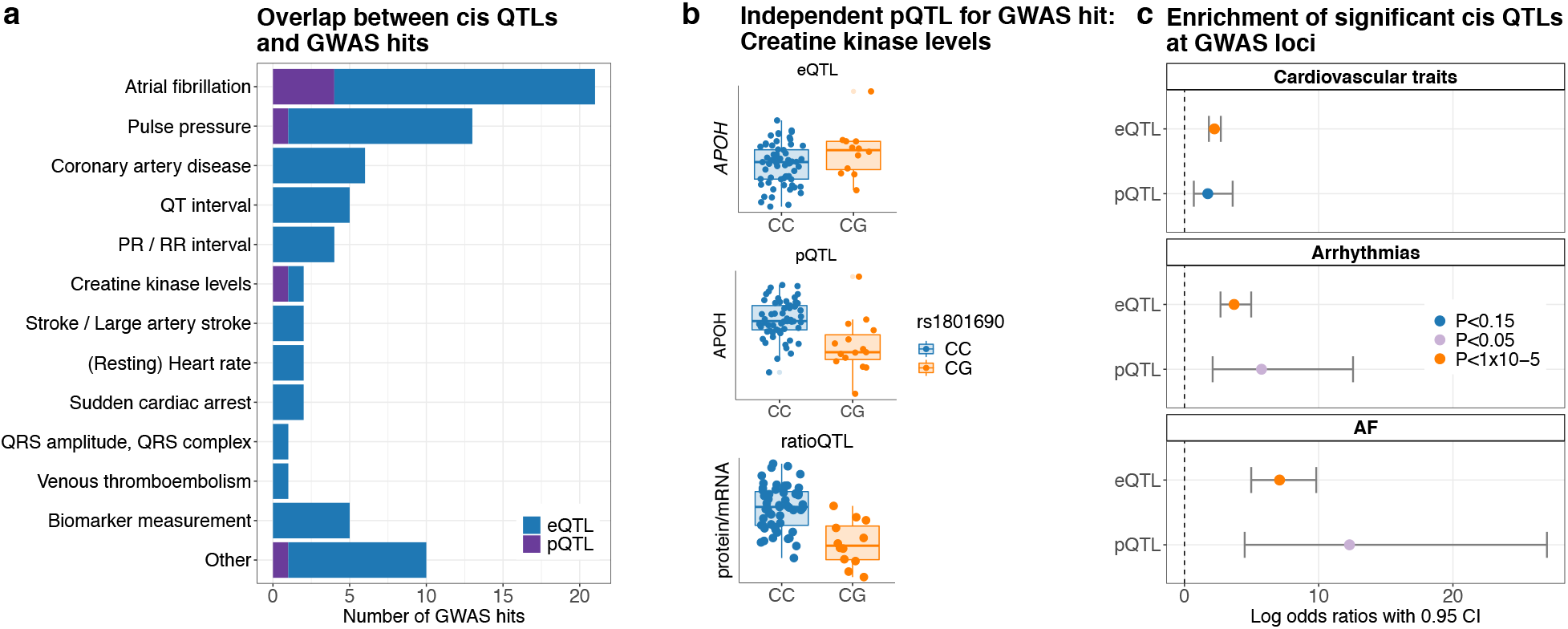
Overlap of cis QTL associations with GWAS hits annotated in the GWAS catalogue. a: Overview of significant cis eQTLs and pQTLs (FDR<0.05) overlapping with GWAS hits for different disease traits. b: Independent pQTL for GWAS hit creatine kinase levels. Shown are the significant cis eQTL, pQTL and ratio QTL for the SNP rs1801690 and the gene APOH (FDR<0.05). In the boxplots, the lower and upper hinges correspond to the first and third quartiles (the 25th and 75th percentiles). The median is denoted by the central line in the box. The upper/lower whisker extends from the hinge to the largest/smallest value no further than 1.5·IQR (interquartile range) from the hinge. c: For three different trait categories (cardiovascular traits, arrhythmias and atrial fibrillation) the enrichment of GWAS hits in significant cis QTLs (FDR<0.05) was evaluated. Enrichments were calculated using Fisher’s exact test (two-sided). QTL, quantitative trait loci; eQTL, expression quantitative trait loci; pQTL, protein quantitative trait loci; ratioQTL, ratio quantitative trait loci; CI, confidence interval;

Of all the overlaps between GWAS hits and cis QTLs (Suppl Table S4), AF-related loci were most abundant (17 eQTLs, four also with pQTL, see Suppl Table S5). Furthermore, we found an independent pQTL overlapping with the GWAS hit for creatine kinase levels (Figure 3b). This genetic effect was not detected on mRNA level illustrating the importance of proteomics data. In addition, we systematically assessed whether significant QTLs are enriched at GWAS loci in the hierarchical groups cardiovascular traits, arrhythmias and AF (two-sided Fisher’s exact test). We identified a strong significant overrepresentation (P<1 × 10^−5^) of eQTLs at GWAS hits for all three groups, and a significant overrepresentation (P<0.05) of pQTLs in variants annotated with arrhythmias and AF (Figure 3c). We showed widespread effects of cis-acting variants on gene expression and protein abundance in atrial tissue and a possible relation to AF.

### Trans QTL analysis

We further extended cis-regulatory analyses by investigating trans effects. Specifically, we addressed a key hypothesis of the omnigenic model,^[10]^ which postulates the existence of core genes. Core genes are central genes with trans associations to AF GWAS loci, whose expression levels directly affect the disease phenotype. Here we sought to identify candidate core genes for AF to understand the contribution of trans-genetic effects in the pathology of AF. To prioritize genes satisfying the properties predicted by the omnigenic model, we evaluated the accumulation of trans effects, their relevance in gene regulatory networks, and the association with AF by the following strategy (Figure 4):

1. We evaluated the cumulated trans effects of AF-associated variants on expression by ranking genes based on their correlation of mRNA and protein abundance with the PRS^[7]^ for AF (eQTS, pQTS).^[12]^ Here, the PRS served as a proxy for an aggregation of AF-related trans effects across the whole genome.
2. To identify genes sharing molecular function and representing biological networks that propagate trans effects to core genes, pathway enrichment analysis (GSEA)^[25, 26]^ was performed on the eQTS and pQTS rankings. Genes driving the enrichment of multiple gene sets were selected as core gene candidates.
3. The link between the core gene candidates and AF was established based on a significant trans eQTL or pQTL for an AF GWAS hit and further supported by differential protein abundance analysis.

**Figure 4:**
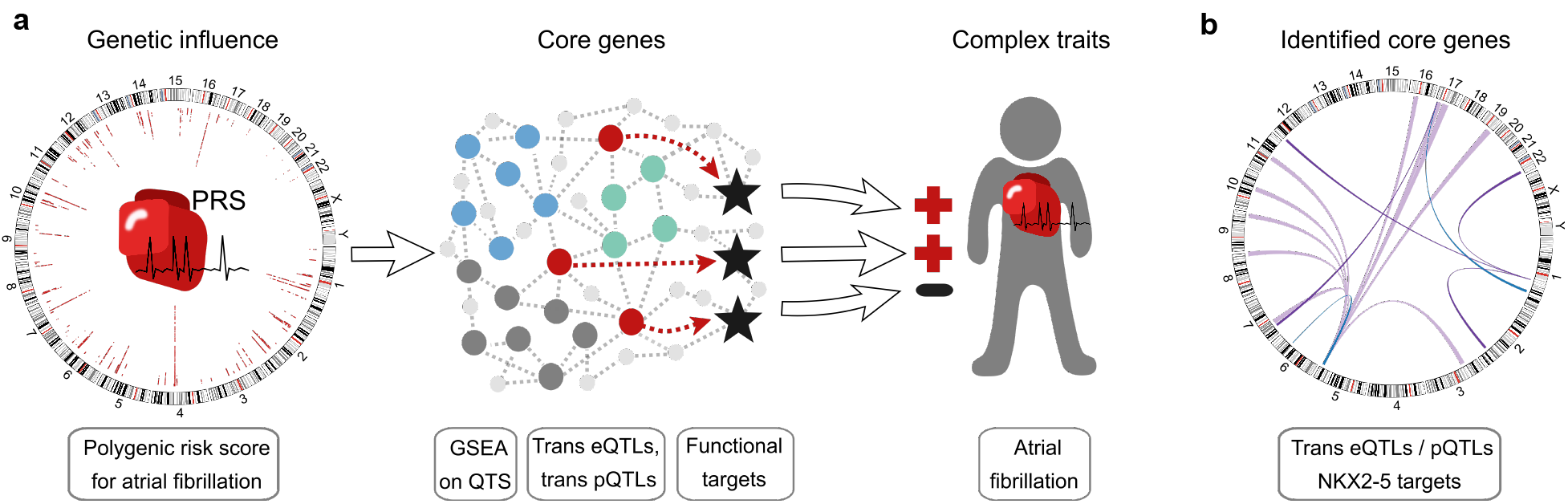
Graphical illustration of the strategy for trans QTL analysis to identify AF-relevant genes. a: Overview: Based on patient-specific PRS values for AF correlated with transcript and protein expression, we performed GSEA to preselect genes for trans eQTL and pQTL analyses from the leading edge of enriched pathways. Core genes were identified as significant trans eQTLs or trans pQTLs. We further assessed their functional targets to investigate the genotype-phenotype relationship in the context of AF. b: Identified core genes as trans eQTLs (blue), trans pQTLs (purple) (FDR<0.2) and functional NKX2-5 targets (light purple). FDR, false discovery rate; AF, atrial fibrillation; GSEA, gene set enrichment analysis; PRS, genome-wide polygenic risk score; QTL, quantitative trait loci for mRNA (eQTL) or protein abundance (pQTL); blue, green or gray dots=core gene candidates; red dots=core genes with trans QTL; stars=functional targets of core genes;

### Identification of putative AF core genes

Based on this strategy, we first used the GO biological process gene set annotations, which are not a priori disease related, to recover processes functionally related to AF. Using all measured transcripts as background for the gene set enrichment, 97 GO biological processes were enriched (adjusted P value <0.05, Suppl Table S6) mostly related to heart muscle or energy metabolism, including the processes *Generation of precursor metabolites and energy*, *Regulation of cardiac muscle contraction*, *Regulation of heart rate*, *Cardiac muscle tissue development*. Restricting the background only to those proteins quantified in our dataset, one GO biological process *Small molecule metabolic process* connected to metabolism was enriched (adjusted P value <0.05, Suppl Table S7).

Our pathway enrichment approach yielded 25 transcripts (Suppl Table S8) and 145 proteins (Suppl Table S9) as core gene candidates that we used to calculate trans QTLs with 109 AF GWAS SNPs. On mRNA level, we identified two trans eQTLs encoding for a cardiac structural protein (rs11658168 - *TNNT2*) and a transcription factor (rs9481842 - *NKX2-5*) (Table 2). On protein level, we discovered four trans pQTLs which are all connected to metabolism (rs11588763 - CYB5R3/NDUFA9/NDUFB3, rs11658168 - HIBADH) (Table 2). Noticeably, three out of four identified genes encode for mitochondrial enzymes (HIBADH) or enzyme subunits (NDUFA9, NDUFB3). Two thirds of the putative core genes have already been mentioned by other studies in the context of arrhythmias and other cardiovascular diseases (detailed findings see Suppl Table S10) which independently replicates the disease link.

**Table 2:** Trans QTL results. Significant trans eQTLs and pQTLs for a FDR<0.2 (Benjamini-Hochberg procedure). Trans analyses were performed on 25 transcripts with 75 samples and 145 proteins with 74 samples for 109 variants associated with atrial fibrillation from the GWAS catalogqt. QTL, quantitative trait loci; FDR, false discovery rate; Mutation known to affect cardiovascular phenotypes; **Mutation known to affect arrhythmias; +Differential expression functional impairment for cardiovascular phenotypes; ++Differential expression or functional impairment for arrhythmias; For details to disease links in literature see Suppl Table S10;

| Variant |  |  | Gene |  | trans QTL |  |  |  |  |
| --- | --- | --- | --- | --- | --- | --- | --- | --- | --- |
| SNP | Chr | Position | Symbol | Chr | QTL | $\beta$ | T value | P value | FDR |
| rs11658168 | chr17 | 7406134 | *TNNT2* * | chr1 | transcript | -0.517 | -4.27 | $6.43 \times 10^{-5}$ | 0.0882 |
| rs9481842 | chr6 | 118974798 | *NKX2-5* ** | chr5 | transcript | -0.593 | -4.27 | $6.54 \times 10^{-5}$ | 0.0882 |
| rs11588763 | chr1 | 154813584 | *CYB5R3* | chr22 | protein | -0.797 | -4.93 | $6.00 \times 10^{-6}$ | 0.0940 |
| rs11588763 | chr1 | 154813584 | *NDUFA9*++ | chr12 | protein | -0.776 | -4.56 | $2.29 \times 10^{-5}$ | 0.129 |
| rs11588763 | chr1 | 154813584 | *NDUFB3*+ | chr2 | protein | -0.937 | -4.54 | $2.47 \times 10^{-5}$ | 0.129 |
| rs11658168 | chr17 | 7406134 | *HIBADH* | chr7 | protein | -0.514 | -4.42 | $3.90 \times 10^{-5}$ | 0.153 |

### NKX2-5 transcription factor network

In order to get more detailed information about complex molecular mechanisms underlying AF, we further analyzed the TF network of NKX2-5 (see Figure 5a) since the TF has already been described in the context of cardiac development,^[27]^ AF,^[28, 29, 30]^ and congenital heart diseases.^[31]^

**Figure 5:**
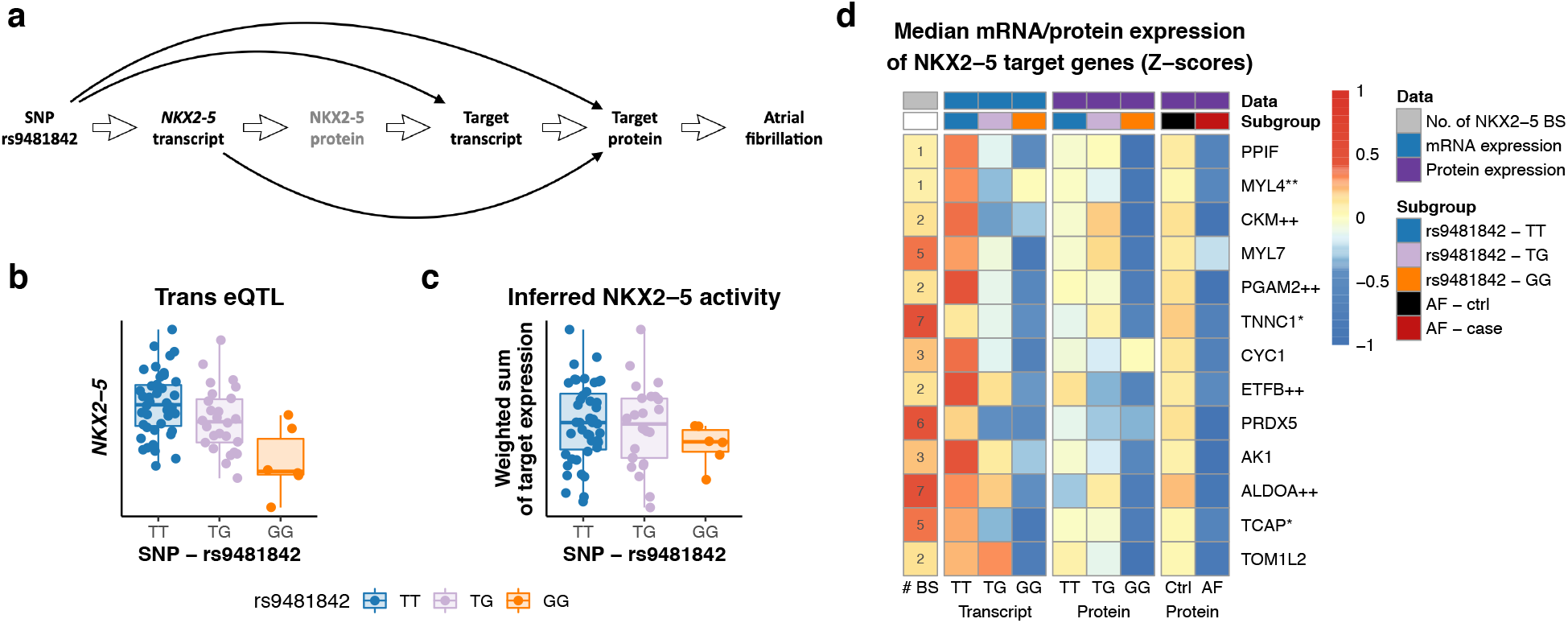
NKX2-5 activity controlled by AF GWAS variant rs9481842. a: Graphical illustration of NKX2-5 TF target gene analysis in AF. b: Strong trans eQTL of the SNP rs9481842 with the *NKX2-5* transcript. c: NKX2-5 activity estimated based on target mRNA expression is still influenced by the rs9481842 genotype. d: Depicted are functional NKX2-5 targets with the number of TF binding sites (column 1), trans eQTL strength (columns 2-4), trans pQTL strength (columns 5-7) and protein level in AF (columns 8-9). The color scale represents median transcript or protein Z-scores per group. AF, atrial fibrillation; QTL, quantitative trait loci; BS, binding site;Mutation known to affect cardiovascular phenotypes; **Mutation known to affect arrhythmias; +Differential expression or functional impairment for cardiovascular phenotypes; ++Differential expression or functional impairment for arrhythmias.

To evaluate the downstream effects of the SNP rs9481842 via the TF NKX2-5 in AF we analyzed the influence of *NKX2-5* transcript levels on target genes by estimating NKX2-5 TF activity. We annotated NKX2-5 binding sites in promoter regions based on published iPSC-derived cardiomyocyte ChIPseq and promoter-capture HiC data to identify almost 10 000 target genes. The number of binding sites per gene in open chromatin regions were counted for each gene and the TF activity (TFA) was computed as the sum of target transcript expression weighted by the number of binding sites. We observed a high correlation between the SNP rs9481842 and the NKX2-5 transcript (cor=−0.43, P=1.4×10^−4^, two-sided Pearson’s correlation, Figure 5b, Suppl Figure S8a) as well as for the direct molecular link between the NKX2-5 transcript and the TF activity (cor=0.36, P=1.3×10^−4^, one-sided Pearson’s correlation, Suppl Figure S8c). In addition, there was a weak association between SNP rs9481842 and the TF activity (cor=−0.13, P=0.145, one-sided Pearson’s correlation, Figure 5b) most likely attributed to the indirect link through NKX2-5. Partial correlation analysis further supported NKX2-5 being the causal link between SNP and target expression (Suppl Table S11).

Next, to elucidate the role of NKX2-5 as a link between the disease variant and AF, we further analyzed its effect on specific targets, which we also prioritized as putative core genes. Overall, we identified 13 functional targets that are significantly influenced by the SNP rs9481842 as well as NKX2-5 transcript levels on both mRNA and protein level (see methods for details, Suppl Table S12). For these 13 targets, we observed a consistent downregulation on mRNA and protein level with respect to the rs9481842 risk allele (Figure 5d). As the core gene model predicts a direct effect of core gene expression on the phenotype,^[10]^ we evaluated the protein abundance of the NKX2-5 target genes in patients with AF compared to patients in sinus rhythm to assess functional connection to the disease. For all targets, AF cases showed lower protein levels (Figure 5d). When adjusting for common risk factors of AF, five out of 13 targets showed a nominal P value smaller 0.05 (Table 3). More importantly, the identified target set collectively displayed a strong association with AF on proteomics level (GSEA P=7.17×10^−5^). This serves as independent validation of the disease link, since these genes were identified based on molecular data in our cohort in combination with public AF annotations without using the actual cohort phenotypes. Furthermore, the majority of the identified proteins are in fact involved in contractile function (MYL4, MYL7, TNNC1, TCAP) or metabolism (PPIF, CKM, AK1, PGAM2, CYC1, ETFB, ALDOA), two mechanisms linked to processes directly influencing AF. At this point, our identified putative core gene point to potential novel targets for further experimental research to better understand molecular consequences of genetics underlying AF.

**Table 3:**
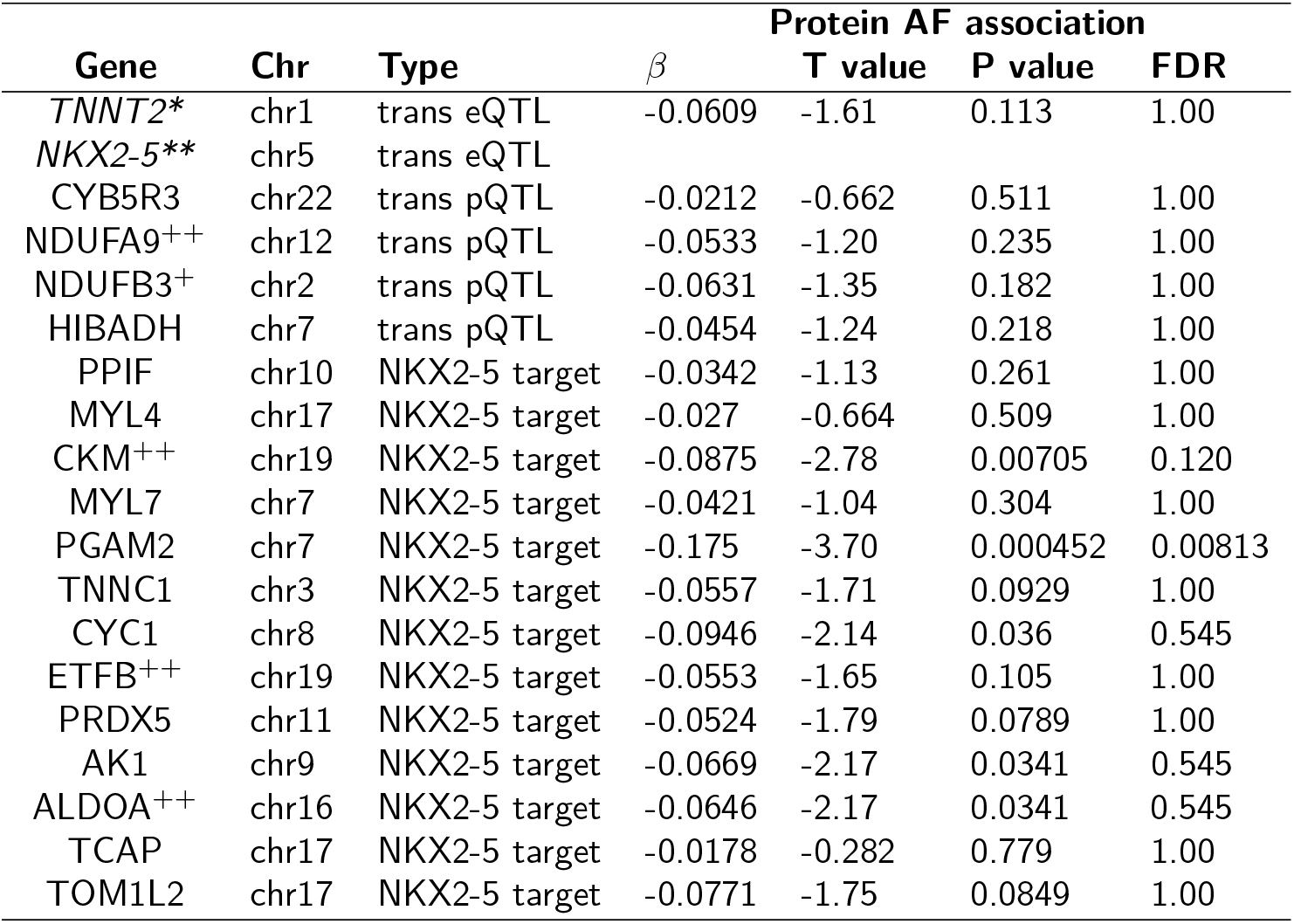
Putative core genes and functional targets with disease association. Proteomics differential abundance results for prevalent AF, corrected for AF-related covariates sex, age, BMI, diabetes, systolic blood pressure, hypertension medication, myocardial infarction, and smoking status (see methods differential protein analysis, N=78, df=66). AF, atrial fibrillation; QTL, quantitative trait loci; FDR, false discovery rate; Mutation known to affect cardiovascular phenotypes; **Mutation known to affect arrhythmias; +Differential expression or functional impairment for cardiovascular phenotypes; ++Differential expression or functional impairment for arrhythmias.

## DISCUSSION

We present a comprehensive multiOMICs analysis that integrates genomics, transcriptomics, and proteomics in human atrial tissue to better understand how genetics are related to molecular mechanisms of AF.

We found widespread genetic effects related to the expression of nearby genes on transcript and protein level. Our integrated eQTL and pQTL analysis allowed the distinction between functional regulatory mechanisms with consequences for mRNA and protein levels. For example, we found many genetic variants exclusively affecting mRNA or protein abundances contributing to a modest overlap between both molecular levels using stringent statistical criteria. Compared to other studies, a similar extent of co-regulation between mRNA and protein levels was previously documented by comparing cis pQTLs in human plasma to cis eQTLs in GTEx tissue^[15]^ (see Suppl “Overlap of eQTLs and pQTLs”, Suppl Table S2). We assume that the use of less stringent significance cutoffs or multiple testing correction as well as more sensitive measurement techniques might achieve a higher overlap as observed by Battle and colleagues for cell-type specific transcriptomics and proteomics in the same lymphoblastoid cell lines.^[14]^ We and others^[14]^ found that proteome-specific pQTLs are enriched in the coding sequence, where post-transcriptional regulatory elements might be affected by sequence variants, which may at least partially explain the divergence between eQTLs and pQTLs. In line with prior studies,^[14, 15, 20]^ we observed large differences in transcript and protein expression as well as their regulation, emphasizing the necessity and benefit of taking multiple molecular entities into account to investigate genotype-phenotype relationships.

To extend the cis QTL analysis, we assessed trans-associations by applying a candidate selection strategy based on the correlation of gene expression with a PRS, a concept termed eQTS.^[12]^ PRS accumulate small genetic effects at many individual genome-wide loci. In the theoretical omnigenic model^[10]^ it has been suggested that these loci are linked to the phenotype by weak trans effects on gene expression, which accumulate in so called core genes. It has been shown that this accumulation of trans effects would lead to strong eQTS associations for core genes.^[12]^ Here we used eQTS and pQTS in combination with gene set enrichment analysis to identify core gene pathways and putative core genes for AF. As core genes are postulated to have trans associations with AF GWAS SNPs we subsequently performed a targeted trans QTL analysis. This strategy allowed the investigation of tissue-specific trans-acting genetic mechanisms underlying AF using a relatively small clinical dataset by reducing the multiple testing burden. The PRS-based gene set enrichment approach revealed cardiac-specific biological pathways associated with the genetic susceptibility for AF which are similar to results identified by Wang and colleagues.^[32]^ We identified different pathways on transcriptome and proteome level which is probably not only caused by biological but also technological reasons like protein coverage. On transcriptome level, the majority of the identified pathways were involved in contractile function and metabolism. In general, these mechanisms have been reported by clinical and experimental studies to play a major role in the pathology of AF.^[33, 34, 35]^ As expected, also the putative core genes identified by trans eQTLs and trans pQTLs were involved in those mechanisms. In addition, all identified transcripts and some of the proteins have been described in the context of arrhythmias or cardiovascular disease (Suppl Table S10). As observed for the cis QTL analysis, the trans analysis revealed similar differences between transcriptomics and proteomics level. Interestingly, none of the trans pQTLs had a significant association on expression level (see Suppl “Trans eQTL / pQTL overlap”, Suppl Table S13). Differences between trans eQTLs and pQTLs for the same gene have previously not been discussed in detail in the literature. Sun and colleagues analyzed overlapping cis QTLs but not trans QTLs.^[15]^ Yao and colleagues analyzed overlaps of cis and trans eQTLs and pQTLs in plasma,^[16]^ however no overlaps were found in trans. Suhre and colleagues validated plasma pQTL findings in other proteomics datasets, but did not evaluate corresponding trans eQTLs.^[13]^ Possible reasons besides the small sample size might be the effect of other genetic variants, post-transcriptional regulation or environmental factors.

To investigate more complex molecular mechanisms underlying AF, we further analyzed the TF network of NKX2-5, since the TF has been described in the context of heart development and arrhythmias. For instance, Benaglio and colleagues discovered that changes in NKX2-5 binding can contribute to electrocardiographic phenotypes.^[30]^ In addition, a NKX2-5 loss-of-function mutation has been reported to be associated with an increased susceptibility to familial AF^[28]^ emphasizing its relevance in the pathology of AF. Although we did not detect NKX2-5 on protein level, we were able to investigate downstream effects by inferring NKX2-5 TF activity based on target transcript expression and functional binding sites. The high correlation between the estimated NKX2-5 TF activity and its mRNA levels shows that NKX2-5 modulates target gene expression. Thus, we also expect altered NKX2-5 protein abundance. These TF activity analyses could only be achieved because of tissue- and cell-type specific annotations, available for NKX2-5. Our analysis suggests that NKX2-5 acts as a transcriptional activator for the majority of genes. The role as activator is consistent with other studies.^[27, 30]^ However, the TF can also function as a transcriptional repressor of genes like ISL1.^[27, 36]^ We were able to detect strong effects of the *NKX2-5* transcript on various target transcript and protein levels. Most of the identified target genes are involved in contractile function or metabolism, two mechanisms highly linked to processes involved in AF. Our unique trans QTL approach revealed direct disease-relevant associations between candidate core genes, NKX2-5 target genes and AF. This finding was confirmed by differential protein abundance between AF patients and controls, which serves as independent validation of the disease relevance of NKX2-5 target genes, since these genes were identified based on molecular data in our cohort in combination with public AF annotations without using the actual cohort phenotypes.

Overall, we successfully integrated multiOMICs data and established a unique approach to investigate not only cis-but also trans-regulatory effects. This provided a platform to generate hypotheses on functional interactions underlying the genetic associations that can be further experimentally investigated.

We acknowledge some limitations that are attributed to common biological and technical factors. First, the use of human heart tissue came with several challenges including restricted sample sizes and heterogeneity of cellular composition. Importantly, changes in cell-type composition and structural remodeling have been described for the pathology of AF.^[37]^ To take differences in the cellular composition into account, we used a fibroblast-score based on a fibroblast-specific gene signature to adjust expression levels in eQTL/pQTL analyses.^[38]^ Second, expression data were generated using microarrays, however, to date, more information can be generated by RNA-seq. Third, human cardiac muscle tissue is dominated by mitochondrial and sarcomere proteins^[39]^ which affects the detection of less abundant proteins such as TFs (e.g. NKX2-5). Therefore, missing TF coverage was not due to data quality but biological and technological restrictions. In addition, only limited functional genomic annotations specific for atrial tissue are currently available including TF-, miRNA- and RBP binding sites. Therefore, we integrated multiple sources to render functional annotations as reliable and accurate as possible. Replication in an independent dataset was not feasible, since this was the first study investigating pQTLs in atrial tissue.

In this study we suggest a novel, integrative analysis of genomics, transcriptomics and proteomics data of human atrial tissue to identify genome-wide genetic effects on intermediate molecular phenotypes in the context of AF. Our multiOMICs approach permits the identification shared and independent effects of cis-acting variants on transcript expression and protein abundance. Furthermore, we proposed a PRS-guided analysis strategy to successfully investigate complex genetic networks even with a limited sample size. By providing these unique tissue-specific OMICs results as a publicly accessible database in an interactive browser, we hope to extend the availability of valuable resources for hypothesis generation, experimental design and target prioritization.

## Methods

### Data and preprocessing

#### Patient cohort

Patients were enrolled in the ongoing observational cohort study AFHRI-B (Atrial fibrillation in high risk individuals-biopsy). Participants were older than 18 years of age and were scheduled to undergo open heart coronary bypass surgery. For the current analyses, N=118 patients with multiOMICs data were available. Cardiovascular risk factor information was collected by questionnaire and from medical records. Baseline blood samples were aliquoted and stored prior to surgery. Atrial appendage tissue remnants were collected when the extracorporeal circulation was started and shock frozen immediately. Follow-up for AF and other cardiovascular disease outcomes was by questionnaire and medical chart review. The observational cohort study was approved by Ethikkommission Ärztekammer Hamburg (PV3982). The study was performed in compliance with the Declaration of Helsinki. The study enrollment and follow-up procedures were in accordance with institutional guidelines. All participants provided written informed consent. Sex stratification of the results was not possible due to the inherently small number of women in the study (Suppl Table S14). Analyses were adjusted for sex where appropriate.

#### Genotypes

The genotype data was generated using the Affymetrix GeneChips Genome-Wide Human SNP Array 6.0, with quality control (QC) at different levels. Using the Birdseed v2 algorithm, PLINK 1.9 and standard quality control procedures,^[40]^ 749 272 SNP were identified in 83 blood samples with a MAF>0.01, HWE exact test P>1 × 10^−6^ and a call rate >98%. Genotypes were further imputed with IMPUTE2^[41]^ based on the 1000 genomes Phase 3 genotypes^[42, 43]^ (per SNP: confident genotype calls with genotype probability >95%, percentage of confident genotype calls across samples >95%) and filtered for HWE P>1 × 10^−4^ resulting in 5 050 128 SNPs for 83 individuals. For QTL analyses, for SNPs with less than 3 individuals with the homozygous-minor-allele genotype, all samples with homozygous-minor-allele genotype were recoded to heterozygous genotype.

#### PRS for AF

The polygenic risk score was calculated based on the LDpred omnigenic score for AF published by Khera et al.^[7]^ To account for a realistic representation of risk score values across the general population, we calculated risk score values together with unrelated 1000 genomes^[42, 43]^ CEU individuals. Phase 3 1000 genomes genotypes were filtered for variants in the risk score and merged with our AFHRI genotypes, resulting in SNP data for 6 730 540 variants out of 6 730 541 in the score. The PRS per individual was computed using the Plink 1.9 score function, imputing missing variants based on the frequency of the risk allele. From this, percentiles across all 490 individuals (1000 genomes: 407, AFHRI cohort: 83) were further used as PRS values for further analysis.

#### mRNA

The mRNA data was generated from human heart atrial appendage tissue samples obtained during heart bypass surgery. They were frozen in liquid nitrogen and pulverized for further analysis. RNA isolation was performed with subsequent assessment of the RNA integrity number (RIN) for quality determination of the samples. HuGene 2.0 ST Arrays were used with the Affymetrix® GeneChip WT Plus Reagent Kit. The R Bioconductor package oligo^[44]^ was used to create expression sets, perform the background correction and quantile-normalization per sample, as well as log-transform the data. Left atrial appendage tissues and samples with a RIN-score smaller than 6 were excluded, in case of replicates the one with the highest RIN-score was used. The mean of multiple transcript clusters annotated to the same gene symbol was used to derive gene level expression values for 26 376 genes in 102 samples.

### Protein

To measure the protein concentrations of the 102 samples, the tissues were homogenized using a micro dismembrator (Braun, Melsungen, Germany) at 2 600 rpm for 2 minutes in 100*μ*l of 8M urea/2M thiourea (UT). Then homogenates were resuspended in 300*μ*l of UT. Nucleic acid fragmentation was gained by sonication on ice three times for 5s each with nine cycles at 80% energy using a Sonoplus (Bandelin, Berlin, Germany). The homogenates were centrifuged at 16 000 x g for one hour at 4°C. After that, protein concentration was determined by Bradford with BSA as standard (SE). 3*μ*g protein were reduced and alkylated and digested with LysC (1:100) for 3h followed by tryptic digestion overnight both at 37°C. Subsequently peptide solutions were desalted on C18 material (ZipTip). Finally mass spectrometry analysis was performed on a LC-ESI-MS/MS machine (LTQ Orbitrap Velos). The Elucidator workflow was used to extract feature intensity and derive protein intensities by summing of all isotope groups with the same peptide annotation for all peptides annotated to one protein (further parameters: Uniprot_Sprot_human_rel. 2016_05: static modification: carbamidomethylation at Cys, variable modification: oxidation at methionine, 2 missed cleavages, fully tryptic, filtered for peptides with FDR<0.5 % corr. to Peptide Teller probability >0.94 and shared peptides were excluded). Intensities for 1 419 proteins with one or more peptides (877 with 2 or more peptides) were quantified for 96 samples, median-normalized and log10-transformed. The original protein concentration was determined as an important technical covariate and therefore used in further analyses.

### Protein-per-mRNA ratios

mRNA and protein measurements were already per-sample quantile-normalized and log-transformed. Both mRNA and protein measurements were additionally quantile-normalized per gene and the ratio was computed as the difference between protein and transcript values.

### Residuals

Per-sample quantile-normalized, log-transformed mRNA and protein values were used to compute residuals. mRNA residuals were derived as the residuals from a linear model explaining mRNA by protein levels, i.e. mRNA ~ *β*_0_ + *β*_1_ protein + *ε*. Protein residuals were derived as the residuals from a linear model explaining protein by mRNA levels, i.e. protein ~ *β*_0_ + *β*_1_ mRNA + *ε*. Covariates were used for further analyses but not for the calculation of residuals.

#### Correction for cell type composition - fibroblast-score

Tissue samples consist of different cell-type compositions. Samples with more fibroblasts probably contain less cardiomyocytes, one of the functionally most relevant cell-types in primary atrial appendage tissue. We used a fibroblast-score based on the sum of expression values of genes upregulated in fibroblasts compared to cardiomyocytes in rats:^[38]^ ELN, FGF10, FOSB, FCRL2, SCN7A, ARHGAP20, CILP, FRAS1, DCDC2, NRG1, AFAP1L2, ITGBL1, NOV, CLEC3B. Cardiomyocyte specific gene signatures were avoided to prevent interfering effects due to structural remodeling common in AF.

### Annotations

#### Genome annotations

Ensembl BioMart^[45]^ GRCh37.p13 hg19 annotations were used as genome annotations (http://feb2014.archive.ensembl.org/biomart/martview/).

#### GWAS catalogue

GWAS annotations were based on the GWAS catalog^[46]^ (https://www.ebi.ac.uk/gwas/, 2019-11-26). We looked at the traits annotated to cardiovascular measurements (EFO_0004298) and cardiovascular disease (EFO_0000319), further referred to as “cardiovascular traits”. We also distinguished the subcategories “arrhythmias”, i.e. all traits annotated to atrial fibrillation, cardiac arrhythmia, sudden cardiac arrest, supraventricular ectopy, early cardiac repolarization measurement, heart rate, heart rate variability measurement, P wave duration, P wave terminal force measurement, PR interval, PR segment, QRS amplitude, QRS complex, QRS duration, QT interval, R wave amplitude, resting heart rate, RR interval, S wave amplitude or T wave amplitude and “AF”, i.e. all traits annotated to Atrial fibrillation or QT interval based on the EFO-mapping (https://www.ebi.ac.uk/gwas/api/search/downloads/trait_mappings, 2019-11-26).

#### VEP

Ensembl Variant Effect Predictions^[47]^ were downloaded from the Ensembl Biomart GRCh37.p13 based on SNP rs-IDs. The label “Missense” was used to summarize all possible missense consequences of the variant (gained stop codon, a frameshift/amino-acid altering/protein-altering variant, a lost start/stop codon, an inframe insertion/deletion).

#### Chromatin states

Roadmap Epigenomics ChromHMM 15 state model coremarks for human heart right atrial appendage^[48]^ E104_15_coreMarks_dense.bed were used to annotate tissue specific chromatin states.

#### Promoter-capture HiC

Promoter-capture HiC data from human iPSC-derived cardiomyocytes^[49]^ (E-MTAB-6014, capt-CM-replicated-interactions-1kb.bedpe) was used to annotate linked promoter regions.

#### Binding sites

TF BS were based on ChIPseq data from the ReMap TF database^[50]^ (ReMap2018 v1.2, bed) filtered for highly expressed genes (log(TPM+1) ≥ 1) in GTEx atrial appendage tissue. Additionally, NKX2-5 binding sites from human iPSC-derived cardiomyocytes^[30]^ (GSE133833) were used. All TF BS were filtered for a minimal overlap of 25 bp with open chromatin regions, i.e. chromatin states “1_TssA”, “2_TssAFlnk”, “10_TssBiv”, “6_EnhG”, “7_Enh”, “11_BivFlnk”, or “12_EnhBiv”.

Fine mapping for functional NKX2-5 BS was done integrating promoter, promoter-capture HiC, chromatin states and NKX2-5 ChIPseq data. Promoter regions were annotated based on Gencode^[51]^ v31lift37 basic and long noncoding RNA transcript start annotations as well as regions linked to those by promoter-capture HiC. ChIPseq binding sites were further overlapped with those promoter regions and filtered for open chromatin regions (details see provided analysis code).

miRNA BS were based on TargetScan 7.2 default predictions for conserved target sites of conserved miRNA families^[52]^ (bed).

RBP BS were derived based on eCLIP data from HepG2 and K562 cell lines provided by the ENCODE Project Consortium^[53, 54]^ (ENCODE, Suppl Table S15). Peak calling was done using the ENCODE uniform processing pipeline, peaks in the bed-files were further filtered for an enrichment >log2(1), a Fisher P value >−log10(0.05) and overlapping peaks were then merged (details see provided analysis code).

#### Methods

Analyses were performed using R 3.4.1. (r-project.org). Genomics data in R was handled using the Bioconductor packages rtracklayer^[55]^ and GenomicRanges.^[56]^

#### Cis QTL covariates including PEER factors

PEER factors were computed using the R package PEER to account for unknown factors in gene expression data.^[17]^ One to 30 PEER factors without additional covariates, with fibroblast-score only and with age, sex, BMI, disease status and fibroblast-score were used as covariates in the QTL analysis (Suppl Figure S1-S2).

#### Cis QTL computation

QTLs were calculated using the R package MatrixEQTL.^[57]^ A cis range of 1 × 10^6^bp and a linear, additive model for genotype effect were used. Expression quantitative trait loci (eQTL), protein quantitative trait loci (pQTL), expression residual quantitative trait loci (res eQTL), protein residual quantitative trait loci (res pQTL) and protein-per-mRNA ratio quantitative trait loci (ratioQTL) analyses were performed for per-sample quantile-normalized as well as additional per-gene quantile normalized expression values (as previously established by Lappalainen and colleagues^[18]^ and the GTEx consortium^[19]^), each for different sets of covariates as described above.

Normalization and covariate sets were optimized for the highest number of QTL genes (i.e. genes, with at least one significant QTL) detected based on a FDR (Benjamini-Hochberg procedure) smaller 0.05 (Suppl Figure S1). For the final analysis, QTLs were computed using per-sample and per-gene quantile-normalized expression values, using only PEER factors without additional covariates (12 PEER factors for eQTLs, 10 PEER factors for pQTLs, 8 PEER factors for res eQTLs and 12 PEER factors for res pQTLs). For ratioQTLs, 9 PEER factors and the fibroblast-score were used as covariates. All tests were two-sided T tests, with 75 samples and 61 degrees of freedom (df) for eQTLs, 75 samples and 63 df for pQTLs, 66 samples and 56 df for res eQTLs, 66 samples and 52 df for res pQTLs and 66 samples and 54 df for ratioQTLs.

#### Definition of functional QTL categories

Shared eQTL / pQTL were defined as SNP-gene pairs with a significant eQTL (FDR<0.05), pQTL (FDR<0.05) and no res eQTL (FDR>0.05) or res pQTL (FDR>0.05), genetic regulation is observable on transcriptomics and proteomics level and variation corresponding to the SNP influence in one OMIC level can be explained and therefore removed by the variation in the other OMIC level. Independent eQTLs were defined as SNP-gene pairs with a significant eQTL (FDR<0.05) and res eQTL (FDR<0.05) but no pQTL (FDR>0.05) and no res pQTL (FDR>0.05), i.e. regulation of SNP is only affecting transcript levels, not proteins. Also, the res eQTL disappears, if the SNP influences protein levels too much, for example a pQTL that barely missed the significance threshold.

Independent pQTLs were defined as SNP-gene pairs with a significant pQTL (FDR<0.05) and res pQTL (FDR<0.05) but no eQTL (FDR>0.05) and no res eQTL (FDR>0.05), i.e. regulation of SNP is only affecting protein levels, not transcripts, i.e. by post-transcriptional regulation.

#### Enrichment of functional elements

Similar as described by Battle and colleagues,[14]} annotations of the top 5 QTL hits per gene were compared to a background distribution (100 background SNPs per QTL SNP) matched for MAF (difference ≤ 0.05) and distance to TSS (difference ≤ 1000bp). Top QTL SNPs per gene were ranked according to the FDR of pQTLs for shared eQTLs / pQTLs, res eQTLs for independent eQTLs and res pQTLs for independent pQTLs. Odds ratios were computed using Fisher’s exact test (two-sided) on the QTL-by-annotation contingency tables.

#### GWAS overlap and enrichments

To determine the overlap between GWAS hits and cis QTLs, we first annotated all GWAS hits for cardiovascular traits in the GWAS catalog^[46]^ with proxies in high linkage-disequilibrium using SNiPA^[58]^ (EUR population, *R*^2^>0.8) as well as significant QTLs (P<1 × 10^−5^). For each of the original GWAS SNPs, the corresponding proxy-gene pair with the strongest QTL was selected to annotate this GWAS hit.

To assess a general enrichment of GWAS hits in QTLs, for all tested QTL SNPs we constructed the cross tables that a SNP has significant QTL (P<1 × 10^−5^) versus was the SNP (or *R*^2^>0.8 proxy) annotated in the GWAS catalog. These tables were evaluated for eQTLs and pQTLs for each of the groups cardiovascular traits, arrhythmias and AF using a two-sided Fisher’s exact test.

#### PRS correlations / eQTS/pQTS rankings

Transcriptomics and proteomics ranking based on PRS correlations were evaluated using linear regression models with additional covariates age, sex, BMI, systolic blood pressure (sysBP), C-reactive protein (CRP) and N-terminal prohormone of brain natriuretic peptide (NT-proBNP). Additionally, the RIN-score or protein concentration were used as technical covariates. The following models were evaluated:

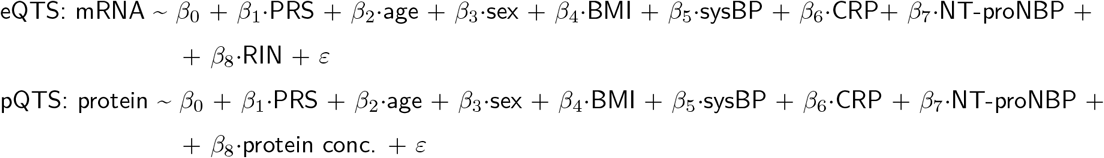

We further used summary statistics (T statistic) for *β*_1_, equivalent to comparing the nested models with/without the PRS and derived corresponding two-sided P values. The degrees of freedom were 66 for mRNA (N=75) and 65 for protein (N=74).

#### Pathway enrichment analysis

Gene set enrichment analysis (GSEA)^[25]^ was performed using the Bioconductor R package fgsea^[26]^ and MSigDB v6.1 gene sets for Gene Ontology biological processes (c5.bp.v6.1.symbols.gmt.txt).^[25, 59, 60]^ To avoid bias towards gene sets specific to human disease as e.g. KEGG pathways, Gene Ontology gene sets, which are not linked to a disease a priori were favoured. The GSEA method was selected to further identify the leading edge genes, which represent the drivers of the enrichment. Enrichments were calculated with 100 000 permutations on eQTS T values (considering gene sets with minimal 15 and maximal 500 transcripts and genes that are highly expressed, i.e. log(TPM+1) ≥ 1, in GTEx atrial appendage tissue) and pQTS T values (considering gene sets with minimal 5 and maximal 500 proteins).

#### SNP and gene candidate selection for trans analyses

To reduce the multiple testing burden, trans analyses were only performed on AF GWAS SNPs and candidate genes derived from the gene set enrichment analysis. We selected all SNPs with MAF ≥ 0.1 that were annotated with atrial fibrillation in the GWAS catalog^[46]^ or the best proxy if the annotated SNP was not measured in our dataset. We further evaluated SNP in high LD using SNiPA^[58]^ (*R*^2^>0.5) and took only the SNP with the highest P value in a recent GWAS,^[3]^ resulting in 109 independent loci. We performed power analysis for the ability to detect strong trans eQTL effects with our fixed sample size (N=75 for eQTLs). The trans eQTL effect size was set to 21.8% of variance explained, which is the strongest trans eQTL found in eQTLGen.^[61]^ In particular we evaluated how many genes can be tested in targeted trans eQTL analysis of all LD pruned AF loci (N=109 SNPs) to still have a power of at least 50% at a Bonferroni adjusted significance level of 5%, based on power calculations for the F test. We found that 26 genes could be tested (Suppl Figure S9). Thus we designed our candidate selection strategy to identify the most promising 26 candidates.

Leading edge genes^[25]^ defined by GSEA on the eQTS / pQTS associations were considered drivers of the enrichments of gene sets. A gene set was considered significantly enriched with a FDR<0.05 (Benjamini-Hochberg procedure). This resulted in 1 539 genes for 97 gene sets on transcript and 145 genes for one gene set on protein level.

Due to the hierarchical structure of the GO biological processes, we favoured genes that were driving the enrichment of multiple gene sets, i.e. also contained in smaller, more specialized child terms. For that reason, we selected all leading edge genes as trans QTL gene candidates that appeared in the transcript leading edge set of 16 or more gene sets, reducing the 1 539 to 25 genes (as based on the power analysis suggesting <26 genes).

Although protein candidates were much more abundant than 26 genes, because of only one significant gene set we could not apply the same selection strategy resulting in no further preselection.

#### Trans QTL computations

Trans QTLs were calculated using the R package MatrixEQTL.^[57]^ An additive linear model was evaluated for 109 SNPs for AF and 25 transcripts as well as 145 proteins. Additional covariates age, sex, BMI, systolic blood pressure, C-reactive protein, N-terminal prohormone of brain natriuretic peptide, the fibro-score and RIN-score/protein concentration for transcripts/proteins were used and resulted in two-sided T test with 75 samples (df=65) for transcripts and 74 samples (df=64). In contrast to the cis QTL analyses, no PEER factors were used as has been previously suggested for trans analyses.^[18]^

#### TF (NKX2-5) target definition selection

We were interested in investigating the link between a GWAS hit to target genes through a trans-eQTL-regulated TF, that was not measured on proteomics level. We therefore investigated the effect of the SNP as well as the TF transcript on target genes on transcriptomics and proteomics level. To do so, we considered only genes with transcriptomics and proteomics measurements and at least one functional TF BS (see methods TF BS). Associations between SNP, transcriptomics and proteomics measurements were evaluated using a linear model with covariates fibroblast-score and RIN-score for transcriptomics measurements and protein concentration for proteomics measurements. We first evaluated four types of associations (two-sided T tests):

1. **Association of GWAS SNP with target transcript (trans eQTL, N=67, df=63):** target transcript *β*_0_ + *β*_1_·SNP + *β*_2_·fibroblast-score + *β*_3_·RIN + *ε*
2. **Independent effects of the SNP on target transcript, that are not mediated by the TF transcript (N=67, df=62):** target transcript *β*_0_ + *β*_1_·SNP + *β*_2_·TF transcript + *β*_3_·fibroblast-score + *β*_4_·RIN + *ε*
3. **Association of target protein with TF transcript (N=79, df=75):** target protein *β*_0_ + *β*_1_·TF transcript + *β*_2_·fibroblast-score + *β*_3_·protein conc. + *ε*
4. **Association of GWAS SNP with target protein (trans pQTL, N=66, df=62):** target protein *β*_0_ + *β*_1_·SNP + *β*_2_·fibroblast-score + *β*_3_·protein conc. + *ε*

Next, we selected only genes with a significant, positive association of the SNP (relative to the allele associated with higher TF expression) with the target transcript (in 1.): *β*_1_<0, P(*β*_1_)<0.05) that vanished, when including the TF transcript in the model (in 2.): P(*β*_1_)>0.2). This was used to ensure that the SNP was not influencing the target directly, but acting through the TF. For the remaining subset, we ranked all targets based on the association of the target protein with the TF transcript and performed FDR correction (Benjamini-Hochberg) on the P values from model (3.) P(*β*_1_) to determine significantly associated genes. Based on this, all targets with FDR<0.05 were defined as functional NKX2-5 targets.

#### Partial correlations

Partial correlations were computed using the R package ppcor.^[62]^

#### Differential proteome analysis for AF

Summary statistics for *β*_1_ (AF) in the following linear model (N=78) were used: protein ~ *β*_0_ + *β*_1_·AF + *β*_2_·sex + *β*_3_·age + *β*_4_·BMI + *β*_5_·diabetes + *β*_6_·sysBP + *β*_7_·hypertension medication + + *β*_8_·myocardial infarction + *β*_9_·smoking status + *β*_10_·fibroblast-score + *β*_11_·protein conc. + *ε*

## Data availability

The data that support the findings of this study are available from the corresponding author upon reasonable request.

## Acknowledgement

This work results from a cooperation in the context with the eMed Junior systems medicine research alliance symAtrial, for which funding by the Federal Ministry of Education and Research was granted to MH (01ZX1408D), MOS (01ZX1408B), RS and TZ (01ZX1408A). MH gratefully acknowledges funding by the Federal Ministry of Education and Research (BMBF, Germany) in the project eMed:confirm (01ZX1708G). RS has received funding from the European Research Council (ERC) under the European Union’s Horizon 2020 research and innovation programme (grant agreement No 648131), from the European Union’s Horizon 2020 research and innovation programme under the grant agreement No 847770 (AFFECT-EU) and German Center for Cardiovascular Research (DZHK e.V.) (81Z1710103). TZ received funding from the eMed Network (01ZX1708A), and from the German Center for Cardiovascular Research (DZHK e.V.) (81X2710170, partner site project; 81X2710105 Shared expertise project). EH was supported financially by the German Center for Cardiovascular Research (DZHK e.V.) (81X2400118 Shared expertise project). Fabian Denbsky setup the prototype of cis QTL processing pipeline during his Master’s thesis. Tim Hartmann assisted with laboratory support, Vishnu Dhople with mass spectrometric data acquisition, and Toray Akcan supplied the processed eCLIP data to derive RBP binding sites. The researchers are indebted to the participants for their willingness to participate in the study and we are very grateful for the staff of the Clinical Cohorts Studies for their important contributions.

## Author contributions

T.Z., R.S. and M.H. conceived the research, supervised the study and obtained funding. J.K., U.V., E.H., T.Z. performed experiments. R.S., C.S.B. and L.C. recruited the patient cohort and collected data. I.A. and M.H. developed computational methods and analyzed the data. B.G. and C.M. contributed to statistical results interpretation. M.O.S. analyzed genotype data. C.M. contributed to the analyses of transcriptomic data. I.A., J.K., T.Z. and M.H. wrote the manuscript with input from all authors. All authors made critical revisions of the manuscript and approved the final version of the manuscript.

## Competing interests

The authors declare no competing interests.

## Notes

### Competing Interest Statement

The authors have declared no competing interest.

### Summary of Updates

Methods and results section updated

