## Supplementary information for "Tissue-specific multiOMICs analysis of atrial fibrillation"

### SUPPLEMENTAL MATERIAL

#### Supplemental Figures and Figure Legends

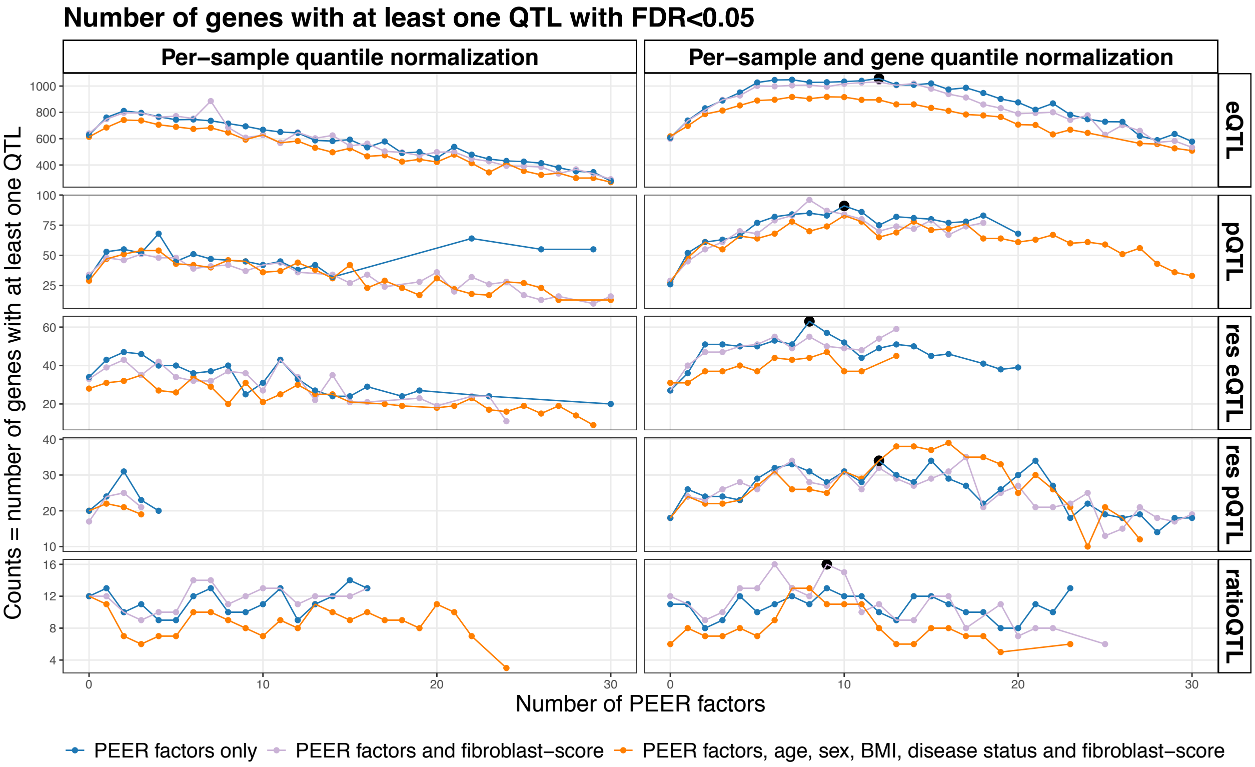


**Figure S1: QTL analysis results for different covariate sets and number of PEER factors.**
PEER analysis was performed to account for unknown variation in the data. Displayed are the number of genes with at least one QTL variant with a FDR<0.05 for different combinations of normalization, number of PEER factors used in the regression and additional covariates. Black dots mark the chosen number of PEER factors and covariates as the maximal number of discovered QTL genes at a FDR<0.05.
FDR, false discovery rate; PEER, probabilistic estimation of expression residuals; eQTL, expression quantitative trait loci; pQTL, protein quantitative trait loci; res eQTL, expression residual quantitative trait loci; res pQTL, protein residual quantitative trait loci; ratioQTL, ratio quantitative trait loci;

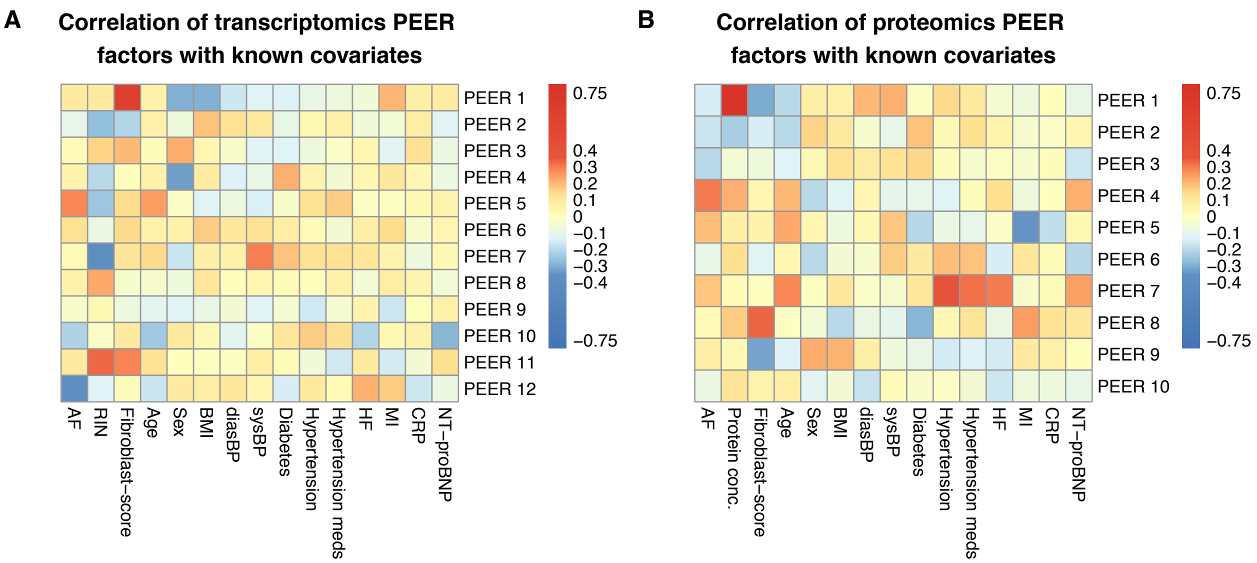


**Figure S2: Correlation of PEER factors with common risk factors of AF and technical covariates.** Pearson correlation of different PEER factors and known risk factors or technical covariates.

A: Transcriptomics analysis. fibroblast-score and RIN-score highly correlate with PEER factors used in the final analysis.

B: Proteomics analysis: fibroblast-score and original sample protein concentration highly correlate with PEER factors used in the final analysis.

PEER, probabilistic estimation of expression residuals; AF, (prevalent) atrial fibrillation; RIN, RNA integrity number; BMI, body mass index; diasBP, diastolic blood pressure; sysBP, systolic blood pressure; HF, heart failure; MI, myocardial infarction; CRP, C-reactive protein; NT-proBNP, N-terminal prohormone of brain natriuretic peptide;

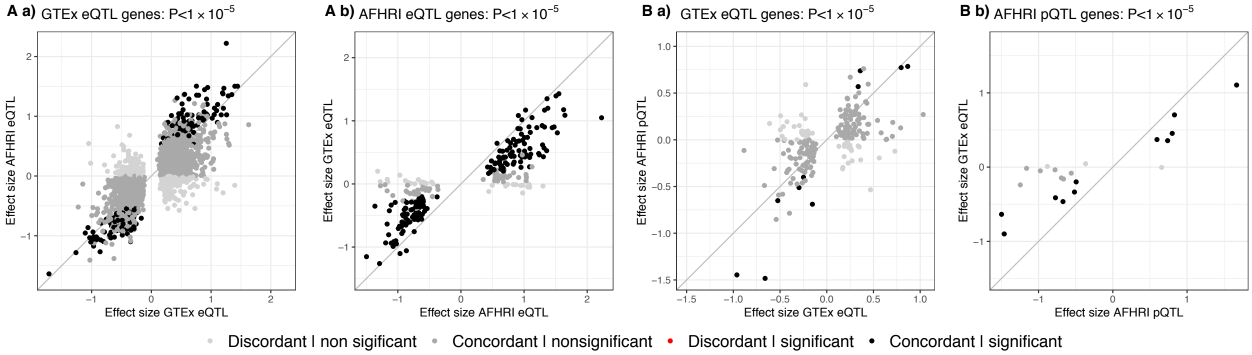


**Figure S3: Comparison of cis eQTL and pQTL results to GTEx cis eQTLs in atrial appendage tissue.**
A: Comparison of effect sizes of eQTLs a) significant eQTLs in GTEx (P$<1 \times 10^{-5}$) and b) significant eQTLs in AFHRI (P$<1 \times 10^{-5}$).
B: Comparison of effect sizes between GTEx eQTLs and AFHRI pQTLs. a) significant eQTLs in GTEx (P$<1 \times 10^{-5}$) and b) significant pQTLs in AFHRI (P$<1 \times 10^{-5}$).
eQTL, expression quantitative trait loci; pQTL, protein quantitative trait loci;

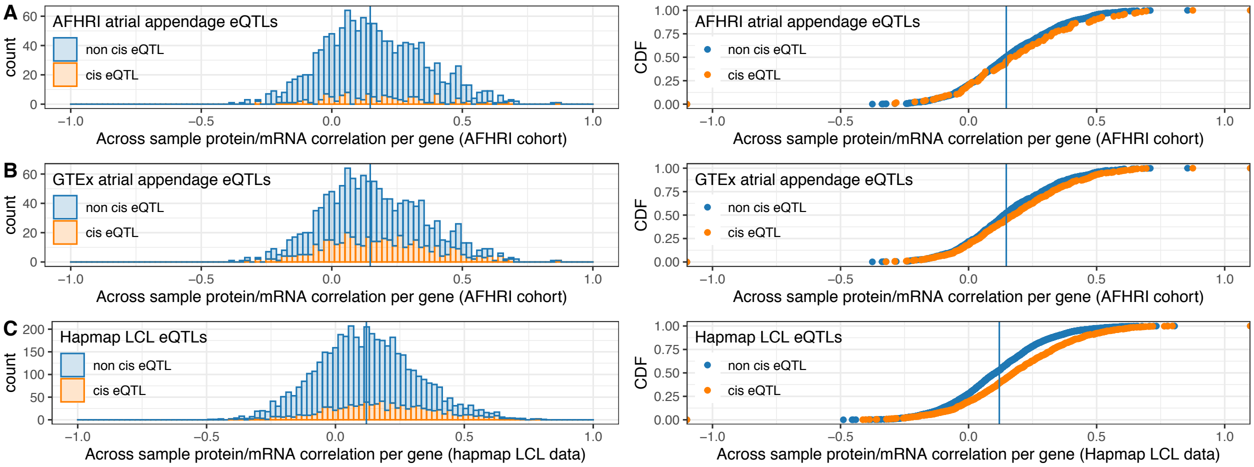


**Figure S4: Pearson correlation between transcript and protein levels dependent on cis eQTL annotations in different datasets.**

A: AFHRI cohort and cis eQTLs. Histogram and cumulative density function.

B: AFHRI cohort and GTEx cis eQTLs annotations. Histogram and cumulative density function.

C: Hapmap LCL data and Hapmap LCL cis eQTL annotations. Histogram and cumulative density function.

eQTL, expression quantitative trait loci; LCL, lymphoblastoid cell line;

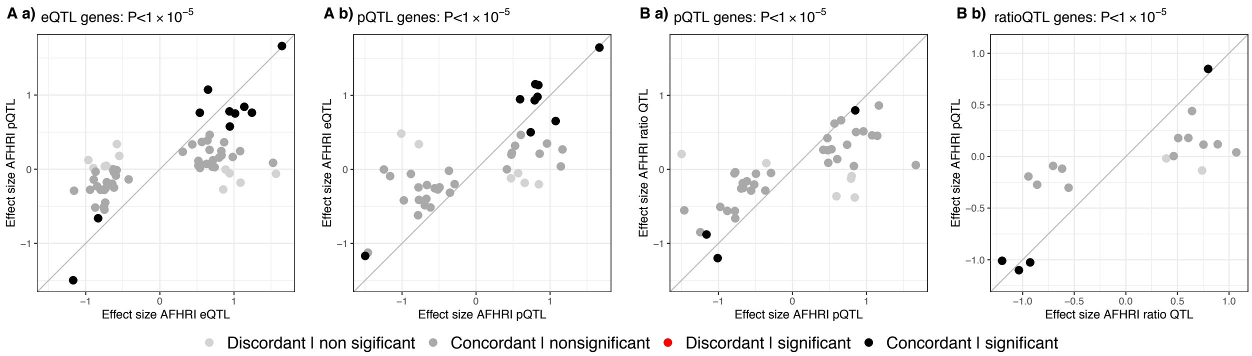


**Figure S5: Between-OMIC comparison of cis QTL results.**
A: comparison of effect sizes of eQTL and pQTL genes for a) significant eQTLs (P$<1 \times 10^{-5}$) and b) significant pQTLs (P$<1 \times 10^{-5}$).
B: comparison of effect sizes of pQTL and ratio QTL genes for a) significant pQTLs (P$<1 \times 10^{-5}$) and b) significant ratio QTLs (P$<1 \times 10^{-5}$).

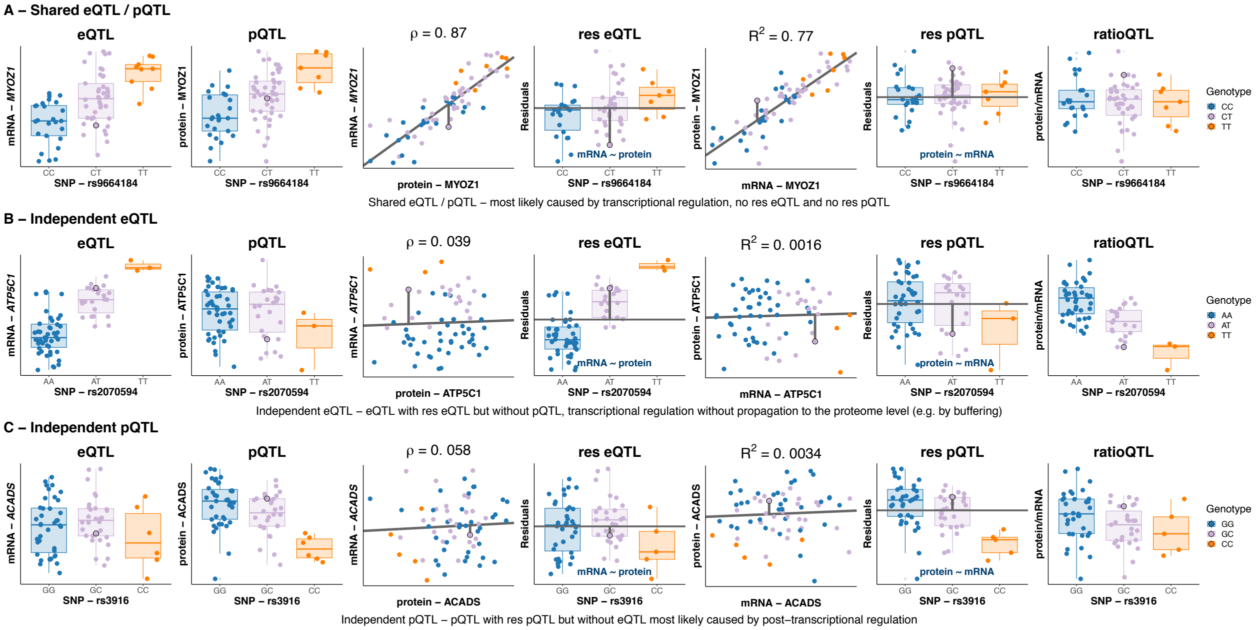


**Figure S6: Extended figure for functional QTL categories.**
QTL boxplots and scatter plots representing residual derivation and mRNA/protein correlation to visualize the three functional QTL categories. In the boxplots, the lower and upper hinges correspond to the first and third quartiles (the 25th and 75th percentiles). The median is denoted by the central line in the box. The upper/lower whisker extends from the hinge to the largest/smallest value no further than 1.5·IQR (interquartile range) from the hinge.

eQTL, expression quantitative trait loci; pQTL, protein quantitative trait loci; res eQTL, expression residual quantitative trait loci; res pQTL, protein residual quantitative trait loci; ratioQTL, ratio quantitative trait loci;

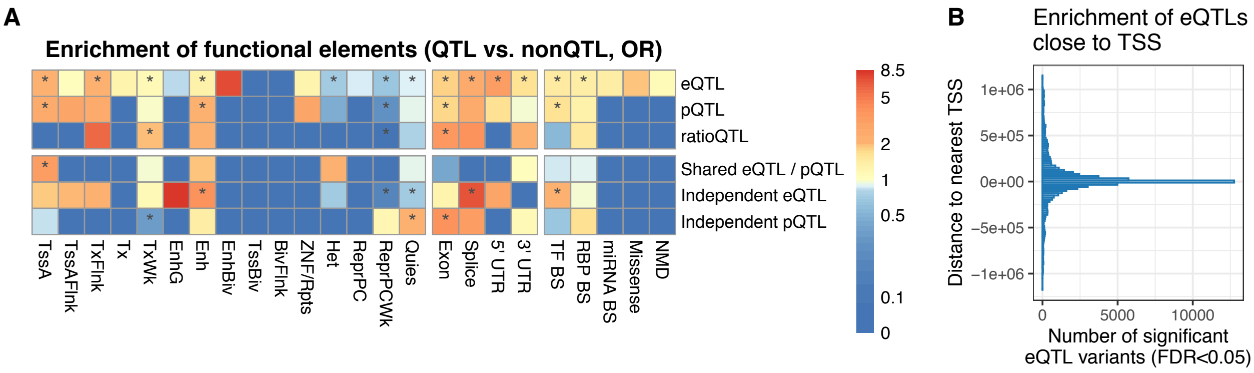


**Figure S7: Enrichment of functional elements for different QTL categories.**
A: Annotations of the top 5 QTL hits per gene were compared to a background distribution (100 background SNPs per QTL SNP) matched for MAF and distance to TSS. Displayed are odds ratios that represent enrichment or depletion, stars mark P values <0.05 for the corresponding Fisher's exact test (two-sided).
B: Enrichment of eQTL hits close to the nearest TSS.
eQTL, expression quantitative trait loci; pQTL, protein quantitative trait loci; ratioQTL, ratio quantitative trait loci; TSS, transcription start site; FDR, false discovery rate;

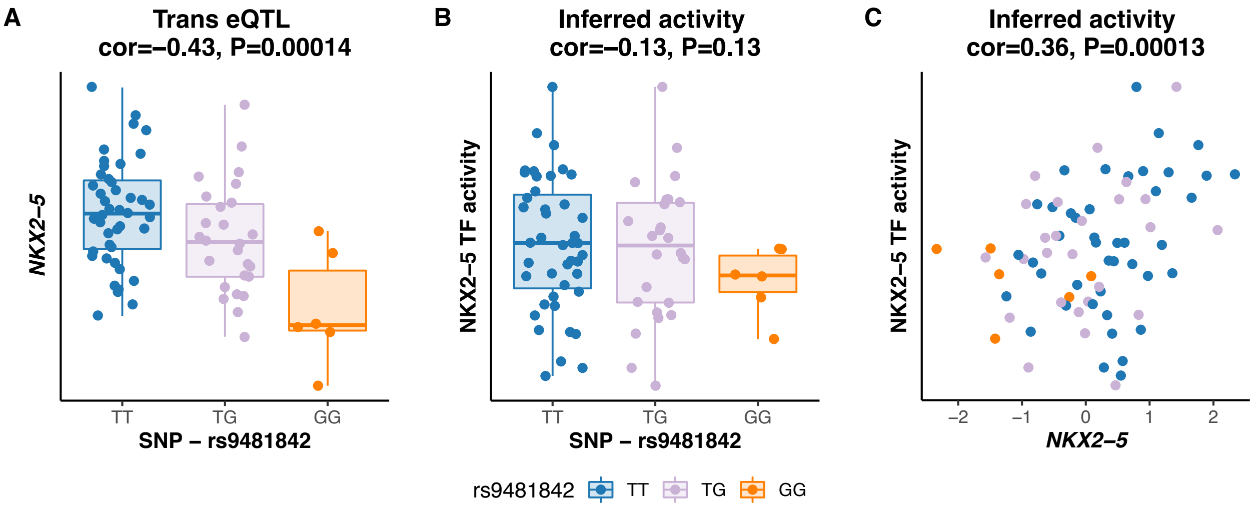


**Figure S8: Causal modeling of NKX2-5 and TF activity.**
A: Trans eQTL rs948182 - *NKX2-5* (two-sided Pearson's correlation).
B: Dependence of TF activity on rs9481842 (one-sided Pearson's correlation).
C: Correlation between *NKX2-5* transcript and TF activity (one-sided Pearson's correlation).
In the boxplots, the lower and upper hinges correspond to the first and third quartiles (the 25th and 75th percentiles). The median is denoted by the central line in the box. The upper/lower whisker extends from the hinge to the largest/smallest value no further than 1.5·IQR (interquartile range) from the hinge.
eQTL, expression quantitative trait loci; TF, transcription factor;

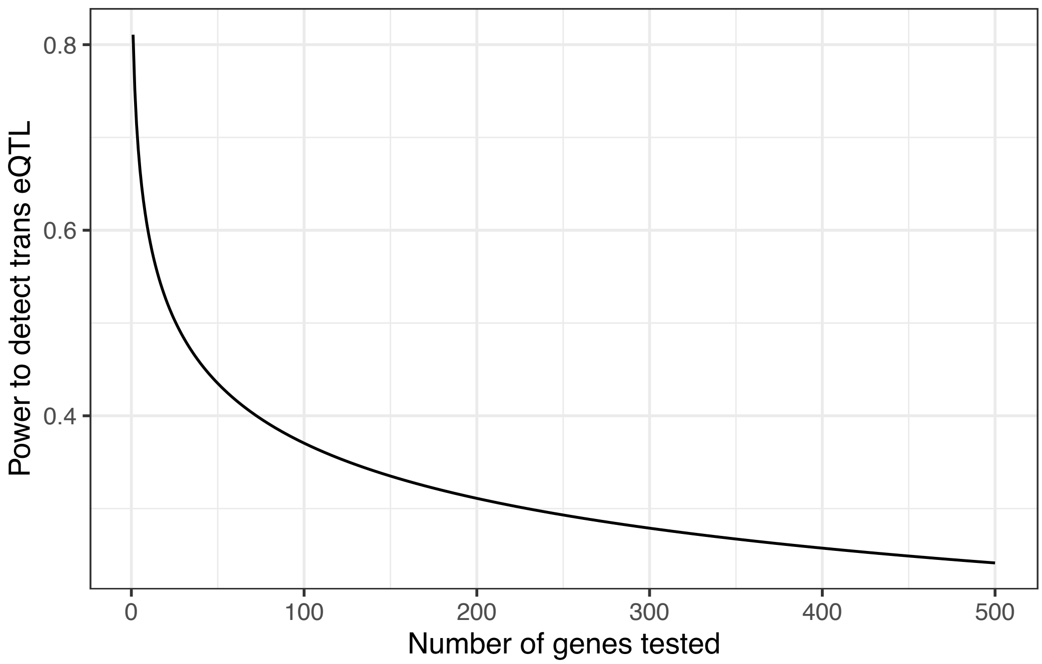


**Figure S9: Trans eQTL power analysis.**
Power analysis for the strongest trans eQTLs (effect size 21.8%), considering 75 samples, 109 SNPs, $\alpha<0.05$ and Bonferroni correction. 26 genes correspond to 50% power.
eQTL, expression quantitative trait loci;

### Supplemental Tables and supporting information

#### Cis QTL results

**Cis QTL replication** We compared QTLs across OMICs for our AFHRI cohort, across studies to the GTEx atrial appendage and across tissues to plasma cis pQTLs. QTLs replicated well across OMICs and studies to the GTEx data, however significance levels varied strongly when compared to plasma pQTL data.
We systematically compared our AFHRI eQTLs to eQTLs for right atrial appendage tissue of the GTEx project. For the comparison, we first filtered the GTEx data for our measured SNPs and genes. We selected the most significant marker for each gene with cis eQTL in our study (P$<1 \times 10^{-5}$) and compared allelic effects. Of all genes available for replication (46%), 66% replicated, 88% showed concordant effects (22% without significance), and none of the eQTLs significant in both datasets showed discordant effects. Using Storey's q-value method,[63] we estimated a replication rate of 83%.
Conversely, of the top GTEx SNPs analyzed in our study, 85% showed concordant allelic effects and 8.5% replicated. The low replication rate is probably due to the large difference in sample size, indicated by the high rate of concordant allelic effects.

**Correlation between mRNA and protein** Protein and transcript levels showed a median correlation of 0.15 per gene and 0.21 per sample. Fitting a linear model to explain protein abundance from transcript expression showed a median $R^2$ of 0.027 per gene and 0.027 per sample (N=79). This general modest mRNA-protein correlation was observed by other comparable datasets as well (Suppl Table S1).[14] Correlations did not increase much if considering only genes with a cis eQTL, however, highly variable genes showed a median correlation of 0.23.
To evaluate the variability of a gene, we compared the logarithm of the variance per gene (over all samples) $\sigma_G$ to the median expression m$_G$ and fit the linear model log($\sigma_G$) ~ $\beta_0 + \beta_1 \cdot$ m$_G$ + $\varepsilon$. Highly variable genes were then defined as all genes $i$ with $\sigma_{G_i} >$ exp($\hat{\beta}_0 + \hat{\beta}_1 \cdot$m$_G$ + 3$\cdot\hat{\sigma}(\hat{\beta}_1)$)

**Table S1: Correlation between mRNA and protein for cis eQTL genes.**
Comparison of median $R^2$ and correlation dependent on existing eQTL annotations. Compared are AFHRI genes non cis eQTL vs. cis eQTL genes, AFHRI genes without an cis eQTL in the GTEx atrial appendage data vs. genes with eQTL in GTEx[19] and Hapmap LCL[14] data without and with an eQTL for the LCL computations.
eQTL, expression quantitative trait score; LCL, lymphoblastoid cell line;

| | AFHRI cohort | | AFHRI cohort | | Hapmap LCL data | |
|---|---|---|---|---|---|---|
| **Measure** | **non cis eQTL** | **cis eQTL** | **non GTEx cis eQTL** | **GTEx cis eQTL** | **LCL non cis eQTL** | **LCL cis eQTL** |
| Median($R^2$) | 0.0265 | 0.0378 | 0.0243 | 0.0354 | 0.0205 | 0.0376 |
| Mean($R^2$) | 0.0634 | 0.0868 | 0.0604 | 0.0771 | 0.0466 | 0.0798 |
| Median(correlation) | 0.145 | 0.174 | 0.137 | 0.175 | 0.105 | 0.176 |
| Mean(correlation) | 0.163 | 0.195 | 0.156 | 0.188 | 0.110 | 0.185 |

**Overlap of eQTL and pQTLs:** Previous studies reported an overlap of 40% of plasma pQTLs[15] with a corresponding eQTL from the GTEx study.[19] However, these eQTLs were not specific for plasma but arose from all available GTEx tissues. Restricting to eQTLs found in whole blood, 19% of plasma pQTLs had a corresponding eQTL. Based on supplemental information given by the authors, the fraction of eQTLs in heart tissues (atrial appendage and left ventricle) that overlapped with plasma pQTLs was much smaller (<7.5%). The other way around, depending on the tissue-specific eQTLs, 12% to 21% of eQTLs also had a plasma pQTL which is well within the range of 17% we observed in our dataset. That implies a qualitatively comparable and reliable overlap between pQTLs and eQTLs which probably is even underestimated. For instance, when using a less stringent significance cutoff of FDR<0.1, a much higher coverage of measured proteins (4 381) and restricting the multiple testing burden by using less SNP-gene pairs in the discovery dataset, Battle and colleagues achieved a much higher replication rate.[14] For matched transcriptomics and proteomics measurements in lymphoblastoid cell lines (LCL), they replicated 35% of eQTL SNP-gene pairs on proteomics level, compared to 14% in our study and 67% of pQTL SNP-gene pairs showed an eQTL, which was significantly more than 16% in our study.
It has to be mentioned, that the way of comparing QTLs differed vastly across studies. Battle and colleagues used

overlapping SNP-gene pairs, without taking linkage information into account.[14] Sun and colleagues first defined lead SNPs in high LD regions ($r^2 \geq 0.8$) for the same gene.[15]
Cis pQTLs in LCLs were also evaluated by Hause and colleagues[20] who found no overlapping cis pQTLs and cis eQTLs at their original significance threshold of FDR<0.05.

**Table S2: Overlap of cis eQTLs and pQTLs in other studies.**
Comparison of cis eQTL/pQTL overlap in previously published studies compared to our dataset. Information about plasma pQTL overlap with GTEx eQTLs is either based on the Sun et al. manuscript (marked with "paper"), or derived from the Sun et al. supplementary table S8 (marked "suppl.").[15]
eQTL, expression quantitative trait loci; pQTL, protein quantitative trait loci; FDR, false discovery rate; LCL, lymphoblastoid cell line;

| Cohort / datasets | Overlap of cis pQTLs with cis eQTL | Overlap of cis eQTLs with cis pQTL |
|---|---|---|
| AFHRI cohort | | |
| QTL genes | n=21 (24%) | n=21 (17%) |
| AFHRI cohort | | |
| SNP-gene pairs | n=642 (16%) | n= 642 (14%) |
| LCL Hause et al.:[20] | none at FDR<0.05 | none at FDR<0.05 |
| LCL Battle et al.:[14] | | |
| SNP-gene pairs | 67% | 35% |
| Plasma pQTL Sun et al.[15] to all GTEx tissues: | | |
| Lead SNPs for same gene, high LD regions ($r^2 \geq 0.8$) | n=224 (40%) (paper) | |
| All sentinel SNPs listed in the Sun et al. supplements | n=320 ( 40%) (suppl.) | |
| Plasma pQTL Sun et al.[15] to GTEx whole blood: | | cis eQTL: P<1.5x10-11 |
| Lead SNPs for same gene, high LD regions ($r^2 \geq 0.8$) | n=117 (19%) (paper) | 12.2% (paper) |
| All sentinel SNPs listed in the Sun et al. supplements | n=152 ( 19%) (suppl.) | |
| Plasma pQTL Sun et al.[15] to GTEx heart tissue: | | |
| Lead SNPs for same gene, high LD regions ($r^2 \geq 0.8$) | | |
| All sentinel SNPs listed in the Sun et al. supplements | n=60 ( 7.5%) (suppl.) | |
| Plasma pQTLSun et al.[15] to GTEx liver: | | cis eQTL: P<1.5x10-11 |
| Lead SNPs for same gene, high LD regions ($r^2 \geq 0.8$) | n=70 () (paper) | 14.8% (paper) |
| All sentinel SNPs listed in the Sun et al. supplements | n=126 ( 16%) (suppl.) | |
| Plasma pQTL Sun et al.[15] to GTEx monocytes: | | cis eQTL: P<1.5x10-11 |
| Lead SNPs for same gene, high LD regions ($r^2 \geq 0.8$) | n=52 () (paper) | 14.7% (paper) |
| All sentinel SNPs listed in the Sun et al. supplements | n=94 ( 12%) (suppl.) | |

**Table S3: Summary of tested data and discovered QTLs.**
Results for a FDR<0.05 (according to Benjamini-Hochberg procedure) and P value $<1 \times 10^{-5}$.
FDR, false discovery rate; eQTL, expression quantitative trait loci; pQTL, protein quantitative trait loci; res eQTL, expression residual quantitative trait loci; res pQTL, protein residual quantitative trait loci; ratioQTL, ratio quantitative trait loci;

| | Tested: | | | FDR<0.05: | | P$<1 \times 10^{-5}$: | |
|---|---|---|---|---|---|---|---|
| | SNPs | Pairs | Genes | Pairs | Genes | Pairs | Genes |
| eQTL | 4 861 118 | 56 139 851 | 16 306 | 57 403 | 1 058 | 40 267 | 552 |
| pQTL | 2 323 504 | 4 508 654 | 1 337 | 4 081 | 91 | 2 543 | 45 |
| ratioQTL | 2 249 758 | 4 198 168 | 1 243 | 563 | 16 | 575 | 18 |
| res eQTL | 2 249 758 | 4 198 168 | 1 243 | 2 261 | 63 | 1 504 | 41 |
| res eQTL | 2 249 758 | 4 198 168 | 1 243 | 1 316 | 34 | 1 194 | 29 |
| Shared eQTL / pQTL | 2 249 758 | 4 198 168 | 1 243 | 430 | 11 | | |
| Independent eQTL | 2 249 758 | 4 198 168 | 1 243 | 1 593 | 37 | | |
| Independent pQTL | 2 249 758 | 4 198 168 | 1 243 | 1 083 | 21 | | |

**Enrichment of functional elements** To further elucidate regulatory mechanisms for the identified QTL associations, we annotated each variant-gene pair with publicly available functional genomics annotations, such as position in the exon, 5' UTR, 3' UTR or splice sites, microRNA binding sites (miRNA BS), transcription factor

binding sites (TF BS), RNA-binding protein binding sites (RBP BS), chromatin states, possible missense mutations or nonsense mediated decay (NMD) (Suppl Figure S7).
Amongst others, there was a significant enrichment (P<0.05, two-sided Fisher's exact test) of active transcription start sites (TssA), enhancer, exon, splice, 5' UTR and 3' UTR regions in eQTLs. Similar results have been reported by Lappalainen and colleagues.[18]
Furthermore, independent pQTLs were enriched in exons, which was previously reported by Battle and colleagues for the comparable group of protein specific QTLs.[14]

#### GWAS catalog overlaps

**Table S4: Cis QTLs overlapping with GWAS hits.**
Number of significant cis QTLs (FDR<0.05) overlapping with variants annotated to cardiovascular traits in the GWAS catalogue (or proxy with $R^2$>0.8).
COPD, Chronic obstructive pulmonary disease; eQTL, expression quantitative trait loci; pQTL, protein quantitative trait loci; ratioQTL, ratio quantitative trait loci; res pQTL, protein residual quantitative trait loci; res eQTL, expression residual quantitative trait loci;

| Trait | eQTL | pQTL | ratioQTL | res pQTL | res eQTL |
|---|---|---|---|---|---|
| Atrial fibrillation | 15 | 3 | 0 | 0 | 0 |
| Pulse pressure | 12 | 1 | 0 | 0 | 0 |
| Coronary artery disease | 5 | 0 | 0 | 0 | 0 |
| QT interval | 5 | 0 | 0 | 0 | 0 |
| Creatine kinase levels | 1 | 1 | 1 | 1 | 0 |
| COPD or resting heart rate (pleiotropy) | 2 | 1 | 0 | 0 | 0 |
| PR interval | 3 | 0 | 0 | 0 | 0 |
| Serum uric acid levels | 2 | 0 | 0 | 0 | 1 |
| Incident atrial fibrillation | 1 | 1 | 0 | 0 | 0 |
| Large artery stroke | 2 | 0 | 0 | 0 | 0 |
| Sudden cardiac arrest | 2 | 0 | 0 | 0 | 0 |
| Age-related disease endophenotypes | 1 | 0 | 0 | 0 | 0 |
| Birdshot chorioretinopathy | 1 | 0 | 0 | 0 | 0 |
| Carotid plaque burden (smoking interaction) | 1 | 0 | 0 | 0 | 0 |
| Circulating myeloperoxidase levels (serum) | 1 | 0 | 0 | 0 | 0 |
| Conotruncal heart defects (inherited effects) | 1 | 0 | 0 | 0 | 0 |
| Coronary artery disease (...) | 1 | 0 | 0 | 0 | 0 |
| Heart rate | 1 | 0 | 0 | 0 | 0 |
| Hematology traits | 1 | 0 | 0 | 0 | 0 |
| Homocysteine levels | 1 | 0 | 0 | 0 | 0 |
| Ischemic stroke | 0 | 0 | 0 | 1 | 0 |
| Left atrial antero-posterior diameter | 1 | 0 | 0 | 0 | 0 |
| Migraine | 1 | 0 | 0 | 0 | 0 |
| Peripheral arterial disease (...) | 1 | 0 | 0 | 0 | 0 |
| Prevalent atrial fibrillation | 1 | 0 | 0 | 0 | 0 |
| QRS complex (Sokolow-Lyon) | 1 | 0 | 0 | 0 | 0 |
| Resting heart rate | 1 | 0 | 0 | 0 | 0 |
| RR interval (heart rate) | 1 | 0 | 0 | 0 | 0 |
| Venous thromboembolism | 1 | 0 | 0 | 0 | 0 |

Due to the strong signal of AF in our QTL results, we further investigated specific hits. We replicated the strong associations at the SYNPO2L locus, where genetic variation strongly influences MYOZ1 but not SYNPO2L. Only 2 (MYOZ1, PCCB) out of the 7 genes with eQTLs that overlapped with AF GWAS hits were measured on proteomics level. The MYOZ1 locus showed a strong eQTL and pQTL, where the QTL and GWAS P values[3] were highly correlated.

**Table S5: Overlap of cis QTLs with GWAS loci for atrial fibrillation.**
QTL hits that overlap with GWAS hits for atrial fibrillation in the GWAS catalogue (or proxy with $R^2 > 0.8$). eQTL, expression quantitative trait loci; pQTL, protein quantitative trait loci;

| Region | QTL variant | Reported variant | Reported gene | QTL gene | QTL type | Pubmed-ID | First author |
|---|---|---|---|---|---|---|---|
| 2p13.3 | rs13028508 | rs10165883 | SNRNP27 | SNRNP27 | eQTL | 29892015 | Roselli C |
| 10q22.2 | rs10824026 | rs10824026 | SYNPO2L | MYOZ1 | eQTL | 29290336 | Nielsen JB |
| 3q22.3 | rs6791611 | rs1278493 | PPP2R3A | PCCB | eQTL | 30061737 | Nielsen JB |
| 5q31.2 | rs9327807 | rs2040862 | WNT8A, NPY6R, MYOT, FAM13B | FAM13B | eQTL | 30061737 | Nielsen JB |
| 2q33.1 | rs159321 | rs295114 | SPATS2L | SPATS2L | eQTL | 29892015 | Roselli C |
| 2q33.1 | rs1347551 | rs3820888 | SPATS2L | SPATS2L | eQTL | 30061737 | Nielsen JB |
| 3q26.33 | rs2339798 | rs4855074 | GNB4 | GNB4 | eQTL | 29892015 | Roselli C |
| 3q26.33 | rs2339798 | rs4855075 | GNB4 | GNB4 | eQTL | 29892015 | Roselli C |
| 1q32.1 | rs951366 | rs4951258 | NUCKS1, SLC41A1 | NUCKS1 | eQTL | 30061737 | Nielsen JB |
| 1q32.1 | rs951366 | rs4951261 | NUCKS1 | NUCKS1 | eQTL | 29892015 | Roselli C |
| 10q22.2 | rs3740293 | rs60212594 | SYNPO2L | MYOZ1 | eQTL | 29892015 | Roselli C |
| 10q22.2 | rs10824026 | rs6480708 | SYNPO2L | MYOZ1 | eQTL | 29892015 | Roselli C |
| 2p13.3 | rs13028508 | rs6546550 | ANXA4/GMCL1 | SNRNP27 | eQTL | 28416818 | Christophersen IE |
| 2p13.3 | rs13028508 | rs6546553 | GMCL1 | SNRNP27 | eQTL | 29892015 | Roselli C |
| 2p13.3 | rs13028508 | rs6747542 | GMCL1, ANXA4 | SNRNP27 | eQTL | 30061737 | Nielsen JB |
| 10q22.2 | rs10824026 | rs7394190 | SYNPO2L | MYOZ1 | eQTL | 28416818 | Christophersen IE |
| 1q32.1 | rs951366 | rs951366 | NUCKS1 | NUCKS1 | eQTL | 29892015 | Roselli C |
| 10q22.2 | rs10824026 | rs10824026 | SYNPO2L | MYOZ1 | pQTL | 28416818 | Christophersen IE |
| 10q22.2 | rs3740293 | rs60212594 | SYNPO2L | MYOZ1 | pQTL | 29892015 | Roselli C |
| 10q22.2 | rs12570126 | rs6480708 | SYNPO2L | MYOZ1 | pQTL | 29892015 | Roselli C |
| 10q22.2 | rs12570126 | rs7394190 | SYNPO2L | MYOZ1 | pQTL | 28416818 | Christophersen IE |

#### Core gene model

#### eQTS / pQTS results with GSEA analysis

**Table S6: Enriched GO terms for eQTS GSEA.**
GSEA results on eQTS T value rankings. GO terms enriched at FDR<0.05. Size refers to the number of genes in the gene set after removing those not evaluated for eQTS.
GO, Gene Ontology; eQTS, expression quantitative trait score; NES, normalized enrichment score; FDR, false discovery rate;

| GO ID | GO term | NES | P value | FDR | size |
|---|---|---|---|---|---|
| GO:0006091 | Generation of precursor metabolites and energy | 2.01 | $1.21\times10^{-5}$ | 0.00398 | 226 |
| GO:0003012 | Muscle system process | 1.95 | $1.23\times10^{-5}$ | 0.00398 | 196 |
| GO:0015980 | Energy derivation by oxidation of organic compounds | 2.09 | $1.25\times10^{-5}$ | 0.00398 | 177 |
| GO:0006936 | Muscle contraction | 1.92 | $1.27\times10^{-5}$ | 0.00398 | 160 |
| GO:0008016 | Regulation of heart contraction | 1.87 | $1.28\times10^{-5}$ | 0.00398 | 143 |
| GO:0045333 | Cellular respiration | 2.14 | $1.31\times10^{-5}$ | 0.00398 | 124 |
| GO:0006119 | Oxidative phosphorylation | 2.02 | $1.40\times10^{-5}$ | 0.00398 | 73 |
| GO:0003015 | Heart process | 2.07 | $1.40\times10^{-5}$ | 0.00398 | 71 |
| GO:0097031 | Mitochondrial respiratory chain complex I biogenesis | 2.15 | $1.47\times10^{-5}$ | 0.00398 | 46 |
| GO:0009060 | Aerobic respiration | 2.14 | $1.47\times10^{-5}$ | 0.00398 | 46 |
| GO:1903522 | Regulation of blood circulation | 1.78 | $2.50\times10^{-5}$ | 0.00408 | 178 |
| GO:0009141 | Nucleoside triphosphate metabolic process | 1.78 | $2.51\times10^{-5}$ | 0.00408 | 168 |
| GO:0090257 | Regulation of muscle system process | 1.80 | $2.57\times10^{-5}$ | 0.00408 | 143 |
| GO:0006937 | Regulation of muscle contraction | 1.87 | $2.69\times10^{-5}$ | 0.00408 | 101 |
| GO:0006956 | Complement activation | -2.42 | $2.87\times10^{-5}$ | 0.00408 | 31 |
| GO:0033108 | Mitochondrial respiratory chain complex assembly | 2.06 | $2.88\times10^{-5}$ | 0.00408 | 57 |
| GO:0002673 | Regulation of acute inflammatory response | -2.40 | $2.94\times10^{-5}$ | 0.00408 | 35 |
| GO:0072376 | Protein activation cascade | -2.20 | $3.01\times10^{-5}$ | 0.00408 | 39 |
| GO:0072350 | Tricarboxylic acid metabolic process | 2.11 | $3.04\times10^{-5}$ | 0.00408 | 34 |
| GO:0006959 | Humoral immune response | -2.18 | $3.37\times10^{-5}$ | 0.00408 | 64 |
| GO:0003013 | Circulatory system process | 1.70 | $3.62\times10^{-5}$ | 0.00408 | 228 |
| GO:0072599 | Establishment of protein localization to endoplasmic reticulum | -2.54 | $3.73\times10^{-5}$ | 0.00408 | 89 |
| GO:0000184 | Nuclear-transcribed mRNA catabolic process, nonsense-mediated decay | -2.05 | $3.91\times10^{-5}$ | 0.00408 | 102 |
| GO:0048738 | Cardiac muscle tissue development | 1.81 | $3.99\times10^{-5}$ | 0.00408 | 110 |
| GO:0070972 | Protein localization to endoplasmic reticulum | -2.33 | $4.01\times10^{-5}$ | 0.00408 | 108 |
| GO:0044033 | Multi-organism metabolic process | -1.90 | $4.18\times10^{-5}$ | 0.00408 | 119 |
| GO:0006413 | Translational initiation | -2.13 | $4.24\times10^{-5}$ | 0.00408 | 124 |
| GO:0006612 | Protein targeting to membrane | -2.33 | $4.31\times10^{-5}$ | 0.00408 | 129 |
| GO:0016567 | Protein ubiquitination | 1.56 | $4.38\times10^{-5}$ | 0.00408 | 488 |
| GO:0050727 | Regulation of inflammatory response | -1.97 | $4.70\times10^{-5}$ | 0.00423 | 156 |
| GO:2000257 | Regulation of protein activation cascade | -2.30 | $5.51\times10^{-5}$ | 0.00456 | 23 |
| GO:0090150 | Establishment of protein localization to membrane | -1.86 | $5.61\times10^{-5}$ | 0.00456 | 214 |
| GO:0006954 | Inflammatory response | -1.72 | $5.63\times10^{-5}$ | 0.00456 | 216 |
| GO:0002027 | Regulation of heart rate | 2.04 | $5.74\times10^{-5}$ | 0.00456 | 59 |
| GO:1903034 | Regulation of response to wounding | -1.87 | $5.90\times10^{-5}$ | 0.00456 | 230 |
| GO:0072657 | Protein localization to membrane | -1.59 | $6.96\times10^{-5}$ | 0.00521 | 289 |
| GO:0006942 | Regulation of striated muscle contraction | 2.03 | $7.13\times10^{-5}$ | 0.00521 | 62 |
| GO:0046487 | Glyoxylate metabolic process | 2.07 | $7.90\times10^{-5}$ | 0.00562 | 21 |
| GO:0006941 | Striated muscle contraction | 1.90 | $8.37\times10^{-5}$ | 0.00569 | 73 |
| GO:0002920 | Regulation of humoral immune response | -2.23 | $8.42\times10^{-5}$ | 0.00569 | 26 |
| GO:0061061 | Muscle structure development | 1.58 | $9.27\times10^{-5}$ | 0.00611 | 306 |
| GO:0050818 | Regulation of coagulation | -2.06 | $9.56\times10^{-5}$ | 0.00615 | 51 |
| GO:0022900 | Electron transport chain | 1.85 | $1.10\times10^{-4}$ | 0.00658 | 83 |
| GO:0002455 | Humoral immune response mediated by circulating immunoglobulin | -2.21 | $1.11\times10^{-4}$ | 0.00658 | 24 |
| GO:0043087 | Regulation of GTPase activity | 1.51 | $1.11\times10^{-4}$ | 0.00658 | 440 |
| GO:0016072 | rRNA metabolic process | -1.70 | $1.12\times10^{-4}$ | 0.00658 | 213 |
| GO:0055117 | Regulation of cardiac muscle contraction | 1.92 | $1.45\times10^{-4}$ | 0.00832 | 55 |
| GO:0019724 | B cell mediated immunity | -1.99 | $1.54\times10^{-4}$ | 0.00868 | 44 |
| GO:0002443 | Leukocyte mediated immunity | -1.85 | $1.87\times10^{-4}$ | 0.0103 | 90 |
| GO:0072521 | Purine-containing compound metabolic process | 1.57 | $2.23\times10^{-4}$ | 0.0121 | 276 |
| GO:0003229 | Ventricular cardiac muscle tissue development | 1.93 | $2.43\times10^{-4}$ | 0.0129 | 34 |
| GO:1903317 | Regulation of protein maturation | -1.92 | $2.57\times10^{-4}$ | 0.0134 | 53 |
| GO:0010575 | Positive regulation of vascular endothelial growth factor production | -2.11 | $2.64\times10^{-4}$ | 0.0135 | 16 |
| GO:0044057 | Regulation of system process | 1.56 | $2.79\times10^{-4}$ | 0.0140 | 297 |
| GO:0045137 | Development of primary sexual characteristics | -1.75 | $2.89\times10^{-4}$ | 0.0142 | 115 |
| GO:0010881 | Regulation of cardiac muscle contraction by regulation of the release of sequestered calcium ion | 1.97 | $3.68\times10^{-4}$ | 0.0178 | 18 |
| GO:0060537 | Muscle tissue development | 1.60 | $3.95\times10^{-4}$ | 0.0187 | 192 |
| GO:0061458 | Reproductive system development | -1.53 | $4.21\times10^{-4}$ | 0.0196 | 236 |
| GO:0006605 | Protein targeting | -1.46 | $4.63\times10^{-4}$ | 0.0212 | 326 |
| GO:0009225 | Nucleotide-sugar metabolic process | -2.02 | $5.19\times10^{-4}$ | 0.0234 | 32 |
| GO:0007548 | Sex differentiation | -1.63 | $5.31\times10^{-4}$ | 0.0235 | 136 |
| GO:0010574 | Regulation of vascular endothelial growth factor production | -2.08 | $5.65\times10^{-4}$ | 0.0246 | 19 |
| GO:0014013 | Regulation of gliogenesis | -1.83 | $6.06\times10^{-4}$ | 0.0260 | 52 |
| GO:1903779 | Regulation of cardiac conduction | 1.83 | $6.31\times10^{-4}$ | 0.0266 | 48 |

| GO ID | GO term | NES | P value | FDR | size |
|---|---|---|---|---|---|
| GO:0045087 | Innate immune response | -1.48 | $6.41\times10^{-4}$ | 0.0266 | 297 |
| GO:0009262 | Deoxyribonucleotide metabolic process | -2.02 | $6.55\times10^{-4}$ | 0.0268 | 29 |
| GO:0030534 | Adult behavior | 1.82 | $6.64\times10^{-4}$ | 0.0268 | 56 |
| GO:1903115 | Regulation of actin filament-based movement | 1.89 | $6.95\times10^{-4}$ | 0.0274 | 28 |
| GO:1901657 | Glycosyl compound metabolic process | 1.54 | $6.99\times10^{-4}$ | 0.0274 | 260 |
| GO:0052547 | Regulation of peptidase activity | -1.50 | $7.16\times10^{-4}$ | 0.0275 | 234 |
| GO:0030258 | Lipid modification | 1.63 | $7.36\times10^{-4}$ | 0.0275 | 138 |
| GO:0055006 | Cardiac cell development | 1.84 | $7.44\times10^{-4}$ | 0.0275 | 42 |
| GO:0043062 | Extracellular structure organization | -1.52 | $7.48\times10^{-4}$ | 0.0275 | 196 |
| GO:0019932 | Second-messenger-mediated signaling | 1.70 | $7.53\times10^{-4}$ | 0.0275 | 101 |
| GO:0010882 | Regulation of cardiac muscle contraction by calcium ion signaling | 1.91 | $7.72\times10^{-4}$ | 0.0278 | 22 |
| GO:0031347 | Regulation of defense response | -1.35 | $8.53\times10^{-4}$ | 0.0303 | 456 |
| GO:0070252 | Actin-mediated cell contraction | 1.78 | $8.63\times10^{-4}$ | 0.0303 | 58 |
| GO:0036498 | IRE1-mediated unfolded protein response | -1.83 | $8.86\times10^{-4}$ | 0.0307 | 50 |
| GO:0055008 | Cardiac muscle tissue morphogenesis | 1.83 | $9.13\times10^{-4}$ | 0.0312 | 40 |
| GO:0007098 | Centrosome cycle | 1.84 | $9.31\times10^{-4}$ | 0.0314 | 33 |
| GO:0086001 | Cardiac muscle cell action potential | 1.84 | $9.85\times10^{-4}$ | 0.0329 | 30 |
| GO:0046578 | Regulation of Ras protein signal transduction | 1.62 | $1.02\times10^{-3}$ | 0.0335 | 129 |
| GO:0007189 | Adenylate cyclase-activating G protein-coupled receptor signaling pathway | 1.89 | $1.11\times10^{-3}$ | 0.0360 | 21 |
| GO:0007507 | Heart development | 1.46 | $1.18\times10^{-3}$ | 0.0381 | 350 |
| GO:0019933 | cAMP-mediated signaling | 1.87 | $1.20\times10^{-3}$ | 0.0381 | 22 |
| GO:0009607 | Response to biotic stimulus | -1.33 | $1.25\times10^{-3}$ | 0.0392 | 477 |
| GO:0060271 | Cilium morphogenesis | -1.62 | $1.26\times10^{-3}$ | 0.0392 | 121 |
| GO:0098901 | Regulation of cardiac muscle cell action potential | 1.86 | $1.31\times10^{-3}$ | 0.0403 | 18 |
| GO:0051298 | Centrosome duplication | 1.86 | $1.38\times10^{-3}$ | 0.0417 | 24 |
| GO:0032940 | Secretion by cell | -1.44 | $1.39\times10^{-3}$ | 0.0417 | 286 |
| GO:0086003 | Cardiac muscle cell contraction | 1.85 | $1.46\times10^{-3}$ | 0.0430 | 24 |
| GO:0055001 | Muscle cell development | 1.66 | $1.46\times10^{-3}$ | 0.0430 | 94 |
| GO:0010880 | Regulation of release of sequestered calcium ion into cytosol by sarcoplasmic reticulum | 1.85 | $1.53\times10^{-3}$ | 0.0446 | 24 |
| GO:0030048 | Actin filament-based movement | 1.70 | $1.66\times10^{-3}$ | 0.0476 | 75 |
| GO:0001501 | Skeletal system development | -1.44 | $1.71\times10^{-3}$ | 0.0486 | 267 |
| GO:0042787 | Protein ubiquitination involved in ubiquitin-dependent protein catabolic process | 1.62 | $1.75\times10^{-3}$ | 0.0492 | 107 |
| GO:0050729 | Positive regulation of inflammatory response | -1.77 | $1.79\times10^{-3}$ | 0.0498 | 48 |

**Table S7: Enriched GO term for pQTS GSEA.**
GSEA results on pQTS T value rankings. GO term enriched at FDR<0.05. Size refers to the number of genes in the gene set after removing those not evaluated for pQTS.
GO, Gene Ontology; pQTS, protein quantitative trait score; NES, normalized enrichment score; FDR, false discovery rate;

| GO ID | GO term | NES | P value | FDR | size |
|---|---|---|---|---|---|
| GO:0044281 | Small molecule metabolic process | -1.68 | $1.27\times10^{-5}$ | 0.0255 | 332 |

**Table S8: Transcriptomics core gene candidates extracted from eQTS GSEA leading edge.**
Genes that appeared in 16 or more GSEA (on eQTS) leading edges for enriched GO terms (FDR<0.05).
#, number of times that the gene appeared in the leading edge;

| Gene | # | Gene | # | Gene | # | Gene | # | Gene | # |
|---|---|---|---|---|---|---|---|---|---|
| RYR2 | 30 | CAV3 | 24 | MYL2 | 21 | TAZ | 19 | TNNT2 | 18 |
| ANK2 | 27 | ATP1A2 | 23 | NKX2-5 | 21 | TNNI3 | 19 | SERPING1 | 17 |
| PKP2 | 25 | GJA5 | 23 | PLN | 20 | CALM1 | 18 | AGT | 16 |
| SLC8A1 | 25 | MYH7 | 23 | C3 | 19 | CALM3 | 18 | C4A | 16 |
| CACNA1C | 24 | DMD | 22 | SCN5A | 19 | MYBPC3 | 18 | MTOR | 16 |

**Table S9: Proteomics core gene candidates extracted from pQTS GSEA leading edge.**
Genes that appeared in the leading edge of the significantly enriched pQTS gene set (FDR<0.05).
#, number of times that the gene appeared in the leading edge;

| Gene | Gene | Gene | Gene | Gene | Gene | Gene |
|---|---|---|---|---|---|---|
| ABHD10 | APOC1 | COX4I1 | GCDH | ME2 | NDUFS8 | PGM5 |
| ACAD10 | APOD | COX5A | GLUD1 | ME3 | NDUFV1 | PPA1 |
| ACADS | AQP1 | COX6C | GOT1 | MPST | NME1 | PRELP |
| ACADVL | ATP1B1 | CRAT | GOT2 | MSRA | NT5C | PRKAR2B |
| ACO1 | ATP5A1 | CS | GPD1L | MTAP | OGDH | PTGES2 |
| ACO2 | ATP5D | CYB5R3 | GPI | MTHFD1 | OGN | PTGIS |
| ACSBG2 | ATP5F1 | CYGB | GSS | NAMPT | OLA1 | QDPR |
| ACSF2 | ATP5J2 | DBT | GSTO1 | NDUFA10 | OXCT1 | SDHA |
| ADA | BCAT2 | DCN | HIBADH | NDUFA5 | P4HB | SDHB |
| ADI1 | BDH1 | DCXR | HIBCH | NDUFA7 | PAFAH1B1 | SDHD |
| ADK | BGN | DDAH2 | HK1 | NDUFA9 | PAM | STOML2 |
| AHCY | BRP44 | ECH1 | HPRT1 | NDUFB11 | PCCB | SUCLA2 |
| AK4 | C3 | ECHS1 | HSD17B10 | NDUFB2 | PCYOX1 | SUCLG1 |
| AKR1A1 | CA1 | ECI2 | HSPA1A | NDUFB3 | PDE5A | TALDO1 |
| AKR7A2 | CA3 | ENO1 | IDH2 | NDUFB8 | PDHA1 | TXNL1 |
| ALDH1A1 | CBR1 | ENO3 | IVD | NDUFB9 | PDXK | UQCR10 |
| ALDH1A2 | CKM | ERLIN2 | LDHA | NDUFC2 | PEPD | UQCRC2 |
| ALDH2 | COQ3 | ETFA | LHPP | NDUFS3 | PGD | UQCRFS1 |
| ALDH4A1 | COQ5 | FH | LUM | NDUFS4 | PGK1 | VCAN |
| ALDH6A1 | COQ7 | FMOD | MCCC1 | NDUFS5 | PGM1 | |
| APOBEC2 | COQ9 | GBAS | MCCC2 | NDUFS7 | PGM2 | |

#### Disease annotations for putative core genes and functional targets evidence in literature

**Table S10: Disease annotations for putative core genes and functional targets in literature.**
Findings relating putative core genes and functional targets to cardiovascular phenotypes.
AF, atrial fibrillation; DCM, Dilated cardiomyopathy; HCM, Hypertrophic cardiomyopathy; Mutation known to affect cardiovascular phenotypes; **Mutation known to affect arrhythmias; $^{+}$Differential expression or functional impairment for cardiovascular phenotypes; $^{++}$Differential expression or functional impairment for arrhythmias.

| Gene | Finding | Clinical phenotypes | First Author |
|---|---|---|---|
| TNNT2* | TNNT2 mutations | DCM | Hershberger[64] |
| NKX2-5** | NKX2-5 mutations including loss of function mutation | Arrhythmia, AF | Jhaveri,[29] Huang[28] |
| NDUFA9$^{++}$ | Impaired complex I function (human atrial tissue) | AF, diabetes | Kanaan[65] |
| NDUFB3$^{+}$ | NDUFB3 deficiency | Cardiomyopathy | El-Hattab[66] |
| MYL4** | Mutation in MYL4 | Familial AF | Orr[67] |
| CKM$^{++}$ | Lower CKM protein expression (human atrial tissue) | AF | Tu[68] |
| | Lower CKM protein expression (human myocardial tissue) | HCM | Coats[69] (Coats Suppl. Table 3) |
| PGAM2$^{++}$ | Lower PGAM2 protein expression (human atrial tissue) | AF | Tu[68] |
| | Lower PGAM2 protein expression (human myocardial tissue) | HCM | Coats[69] (Coats Suppl. Table 3) |
| TNNC1* | Mutation in TNNC1 | Cardiomyopathy | Parvatiyar[70] |
| ETFB$^{++}$ | Lower ETFB protein expression (human atrial tissue) | AF | Tu[68] |
| | ETFB mutation | Arrhythmias, HCM, DCM, conduction defects | Florian[71] |
| ALDOA$^{++}$ | Lower ALDOA protein expression (human atrial tissue) | AF | Tu[68] |
| | Lower ALDOA protein expression (human myocardial tissue) | HCM | Coats[69] (Coats Suppl. Table 3) |
| TCAP* | TCAP gene mutations | HCM, DCM | Hayashi[72] |
| TOM1L2 | Differentially expressed | AF together with neurocognitive decline | Dalal[73] |

Additionally, as stated by Wang and colleagues,[32] multiple genes were either identified by their integrative omics approach or mentioned in the OMIM[74] database.

#### NKX2-5 causal modeling

**Table S11: Partial correlation analysis of *NKX2-5* expression linking the SNP rs9481842 and TF activity.**
Two-sided Pearson's correlation tests unless stated otherwise.
TF, transcription factor; *one-sided Pearson's correlation test;

| Measure | SNP and *NKX2-5* mRNA | SNP and TF activity | *NKX2-5* mRNA and TF activity |
|---|---|---|---|
| Correlation | -0.43 (P = $1.4 \times 10^{-4}$) | -0.13 (P = 0.13)* | 0.36 (P = $1.3 \times 10^{-4}$)* |
| Partial correlation | -0.41 (P = $3 \times 10^{-4}$) | 0.007 (P = 0.95) | 0.3 (P = 0.011) |
| Condition | TF activity | *NKX2-5* mRNA | SNP |

**Table S12: NKX2-5 target correlations with trans eQTL SNP rs9481842 and *NKX2-5* transcript as well as AF disease association.**
Associations were computed using a linear model with covariates (fibroblast-score, RIN-score / sample protein concentration). Correlation of SNP rs9481842 with target transcript (trans eQTL) and target protein (trans pQTL) were evaluated as well as the correlation between *NKX2-5* transcript expression and target protein expression. Proteomics differential expression results included common risk factors for AF as covariates (age, sex, BMI, diabetes, systolic blood pressure, hypertension medication, myocardial infarction and smoking).
AF, atrial fibrillation; eQTL, expression quantitative trait loci; pQTL, protein quantitative trait loci; FDR, false discovery rate; Mutation known to affect cardiovascular phenotypes; **Mutation known to affect arrhythmias; $^{+}$Differential expression or functional impairment for cardiovascular phenotypes;$^{++}$Differential expression or functional impairment for arrhythmias.

| ***NKX2-5 target*** | | **SNP rs9481842** | | | | ***NKX2-5* - target protein association** | | | **Disease association protein and AF** | |
|---|---|---|---|---|---|---|---|---|---|---|
| | | **Trans eQTL** | | **Trans pQTL** | | | | | | |
| **Gene** | **No. BS** | $\beta$ | **P value** | $\beta$ | **P value** | $\beta$ | **P value** | **FDR** | $\beta$ | **P value** |
| PPIF | 2 | -0.131 | 0.0081 | -0.0388 | 0.0145 | 0.172 | $2.10\times10^{-4}$ | 0.00824 | -0.0342 | 0.261 |
| MYL4** | 1 | -0.087 | 0.00921 | -0.0359 | 0.0820 | 0.211 | $3.98\times10^{-4}$ | 0.00824 | -0.0270 | 0.509 |
| CKM$^{++}$ | 2 | -0.115 | 0.0101 | -0.0188 | 0.303 | 0.180 | $4.56\times10^{-4}$ | 0.00824 | -0.0875 | 0.00705 |
| MYL7 | 5 | -0.125 | 0.0038 | -0.0298 | 0.177 | 0.208 | $4.74\times10^{-4}$ | 0.00824 | -0.0421 | 0.304 |
| PGAM2$^{++}$ | 2 | -0.214 | 0.00307 | -0.0293 | 0.284 | 0.255 | $7.35\times10^{-4}$ | 0.0107 | -0.175 | 0.000452 |
| TNNC1* | 8 | -0.0699 | 0.0359 | -0.00809 | 0.623 | 0.147 | $1.69\times10^{-3}$ | 0.0211 | -0.0557 | 0.0929 |
| CYC1 | 3 | -0.121 | 0.00278 | -0.00651 | 0.749 | 0.175 | $3.16\times10^{-3}$ | 0.0307 | -0.0946 | 0.0360 |
| ETFB$^{++}$ | 3 | -0.0924 | 0.0110 | -0.0336 | 0.0563 | 0.152 | $3.39\times10^{-3}$ | 0.0307 | -0.0553 | 0.105 |
| PRDX5 | 6 | -0.0710 | 0.00641 | -0.0149 | 0.307 | 0.131 | $3.52\times10^{-3}$ | 0.0307 | -0.0524 | 0.0789 |
| AK1 | 4 | -0.0654 | 0.0268 | -0.0224 | 0.190 | 0.138 | $4.17\times10^{-3}$ | 0.0312 | -0.0669 | 0.0341 |
| ALDOA$^{++}$ | 11 | -0.0555 | 0.0469 | -0.00187 | 0.909 | 0.125 | $5.50\times10^{-3}$ | 0.0368 | -0.0646 | 0.0341 |
| TCAP* | 5 | -0.114 | 0.00707 | -0.0365 | 0.266 | 0.244 | $6.90\times10^{-3}$ | 0.0429 | -0.0178 | 0.779 |
| TOM1L2 | 2 | -0.0886 | 0.0157 | -0.0292 | 0.222 | 0.170 | $8.16\times10^{-3}$ | 0.0473 | -0.0771 | 0.0849 |

#### Trans eQTL / pQTL overlap

**Table S13: Trans QTL results.**
Significant trans eQTLs and pQTLs at a FDR <0.2 (Benjamini-Hochberg procedure). 109 SNPs annotated with atrial fibrillation were tested for association with 25 transcripts and 145 proteins.
QTL, quantitative trait loci; eQTL, expression quantitative trait loci; pQTL, protein quantitative trait loci; FDR, false discovery rate; Mutation known to affect cardiovascular phenotypes; **Mutation known to affect arrhythmias; $^{+}$Differential expression or functional impairment for cardiovascular phenotypes;$^{++}$Differential expression or functional impairment for arrhythmias.

| | Variant | | Gene | | | Trans eQTL | | | | Trans pQTL | | |
|---|---|---|---|---|---|---|---|---|---|---|---|---|
| **SNP** | **Chr** | **Position** | **Transcript** | **Chr** | $\beta$ | **T value** | **P value** | **FDR** | $\beta$ | **T value** | **P value** | **FDR** |
| rs11658168 | chr17 | 7 406 134 | _TNNT2*_ | chr1 | -0.517 | -4.27 | $6.43\times10^{-5}$ | **0.0882** | -0.215 | -1.43 | $1.57\times10^{-1}$ | 0.914 |
| rs9481842 | chr6 | 118 974 798 | _NKX2-5**_ | chr5 | -0.593 | -4.27 | $6.54\times10^{-5}$ | **0.0882** | | | | |
| | **Variant** | | **Gene** | | | **Trans eQTL** | | | | **Trans pQTL** | | |
| **SNP** | **Chr** | **Position** | **Protein** | **Chr** | $\beta$ | **T value** | **P value** | **FDR** | $\beta$ | **T value** | **P value** | **FDR** |
| rs11588763 | chr1 | 154 813 584 | CYB5R3 | chr22 | -0.119 | -0.527 | $6.00\times10^{-1}$ | 0.998 | -0.797 | -4.93 | $6.00\times10^{-6}$ | **0.094** |
| rs11588763 | chr1 | 154 813 584 | NDUFA9$^{++}$ | chr12 | 0.257 | 1.16 | $2.49\times10^{-1}$ | 0.986 | -0.776 | -4.56 | $2.29\times10^{-5}$ | **0.129** |
| rs11588763 | chr1 | 154 813 584 | NDUFB3$^{+}$ | chr2 | 0.29 | 1.31 | $1.94\times10^{-1}$ | 0.981 | -0.937 | -4.54 | $2.47\times10^{-5}$ | **0.129** |
| rs11658168 | chr17 | 7 406 134 | HIBADH | chr7 | -0.144 | -0.861 | $3.93\times10^{-1}$ | 0.998 | -0.514 | -4.42 | $3.90\times10^{-5}$ | **0.153** |

#### AFHRI cohort

**Table S14: AFHRI cohort baseline table.**
AF, atrial fibrillation; PRS, genome-wide polygenic score; BMI, body-mass index; BP, blood pressure;

| Variable | Total (N=118) | Women (N=13) | Men (N=105) |
|---|---|---|---|
| Genotypes measured, n (%) | 83 (70) | 6 (46) | 77 (73) |
| Transcriptomics measured, n (%) | 102 (86) | 10 (77) | 92 (88) |
| Proteomics measured, n (%) | 96 (81) | 8 (61) | 88 (84) |
| All measured, n (%) | 66 (56) | 3 (23) | 63 (60) |
| AF, n (%) | 15 (13) | 1 (8) | 14 (13) |
| AF PRS, median (IQR) | 32.40 (32.33-32.48) | 32.41 (32.40-32.42) | 32.40 (32.33-32.49) |
| Age, median (IQR), y | 66.8 (59.5-73.5) | 67.6 (63.5-71.8) | 66.4 (59.0-73.5) |
| BMI, median (IQR), kg/m$^2$ | 27.8 (24.8-30.4) | 28.3 (24.1-30.1) | 27.8 (25.0-30.4) |
| systolic BP, median (IQR), mmHg | 135 (122-145) | 137 (120-150) | 135 (123-145) |
| diastolic BP, median (IQR), mmHg | 76 (70-82) | 72 (69-80) | 76 (70-82) |
| Hypertension, n (%) | 105 (89) | 13 (100) | 92 (88) |
| Hypertension medication, n (%) | 98 (83) | 13 (100) | 85 (81) |
| Diabetes, n (%) | 36 (31) | 4 (31) | 32 (30) |
| Diabetes medication, n (%) | 33 (28) | 3 (23) | 30 (29) |
| Myocardial infarction, n (%) | 45 (38) | 5 (38) | 40 (38) |
| Smoking, n (%) | 28 (24) | 1 (8) | 27 (26) |
| fibroblast-score, median (IQR) | 80.43 (79.52-81.87) | 80.97 (80.43-81.57) | 80.34 (79.25-81.89) |
| RIN-score, median (IQR) | 7.7 (7.1-8.1) | 7.2 (6.7-7.8) | 7.7 (7.2-8.1) |
| Protein concentration, median (IQR), $\mu g/\mu l$ | 0.87 (0.45-1.31) | 0.81 (0.45-1.35) | 0.87 (0.45-1.30) |

#### Supporting data resources

**Table S15: ENCODE resources used to annotate RNA-binding proteins (RBP) binding sites.**
eCLIP data for all RBPs for Hep2 and K562 cell lines provided by Gene Yeo, UCSD (https://www.encodeproject.org/report/?type=Experiment&status=released&replicates.library.biosample.donor.organism.scientific_name=Homo+sapiens&biosample_ontology.classification=cell+line&assay_title=eCLIP&limit=all).

| Accession | Assay | Target gene | Cell line | Accession | Assay | Target gene | Cell line | Accession | Assay | Target gene | Cell line |
|---|---|---|---|---|---|---|---|---|---|---|---|
| ENCSR999YGP | eCLIP | GRWD1 | K562 | ENCSR648LAH | eCLIP | DDX3X | HepG2 | ENCSR001KKZ | eCLIP | PHF6 | K562 |
| ENCSR249ROI | eCLIP | HNRNPC | K562 | ENCSR384MWO | eCLIP | CSTF2 | HepG2 | ENCSR721HPX | eCLIP | G3BP1 | HepG2 |
| ENCSR820UYE | eCLIP | PABPN1 | HepG2 | ENCSR154CSN | eCLIP | DDX52 | K562 | ENCSR464OSH | eCLIP | FUS | HepG2 |
| ENCSR365NVO | eCLIP | TRA2A | K562 | ENCSR331VNX | eCLIP | FMR1 | K562 | ENCSR906ZJF | eCLIP | SDAD1 | K562 |
| ENCSR337XGI | eCLIP | SAFB | HepG2 | ENCSR987NYS | eCLIP | FAM120A | HepG2 | ENCSR754NDA | eCLIP | RBM15 | HepG2 |
| ENCSR977OXG | eCLIP | PRPF4 | HepG2 | ENCSR861GYE | eCLIP | LIN28B | HepG2 | ENCSR181NRW | eCLIP | ZC3H8 | K562 |
| ENCSR862QCH | eCLIP | U2AF1L5,U2AF1 | K562 | ENCSR970FEW | eCLIP | DDX52 | HepG2 | ENCSR322HHA | eCLIP | TIAL1 | HepG2 |
| ENCSR240MVJ | eCLIP | HNRNPU | HepG2 | ENCSR973HOJ | eCLIP | FXR2 | HepG2 | ENCSR867ZVK | eCLIP | YWHAG | K562 |
| ENCSR529FKI | eCLIP | YBX3 | K562 | ENCSR352STY | eCLIP | SSB | HepG2 | ENCSR584TCR | eCLIP | TARDBP | K562 |
| ENCSR841EQA | eCLIP | TAF15 | HepG2 | ENCSR050BDZ | eCLIP | SDAD1 | HepG2 | ENCSR999WKT | eCLIP | DDX24 | K562 |
| ENCSR022BVV | eCLIP | LSM11 | K562 | ENCSR685AUR | eCLIP | ZNF800 | HepG2 | ENCSR456KXI | eCLIP | LARP7 | K562 |
| ENCSR267OLV | eCLIP | SF3B4 | K562 | ENCSR089BXO | eCLIP | ABCF1 | K562 | ENCSR840DRD | eCLIP | CSTF2T | K562 |
| ENCSR828ZID | eCLIP | HNRNPK | HepG2 | ENCSR145NLR | eCLIP | DDX51 | K562 | ENCSR653HQC | eCLIP | DROSHA | K562 |
| ENCSR406OOZ | eCLIP | SUB1 | HepG2 | ENCSR989VIY | eCLIP | SRSF1 | HepG2 | ENCSR331MIC | eCLIP | SF3A3 | HepG2 |
| ENCSR062NNB | eCLIP | IGF2BP2 | K562 | ENCSR194HZU | eCLIP | NOLC1 | HepG2 | ENCSR993OLA | eCLIP | IGF2BP3 | HepG2 |
| ENCSR267UCX | eCLIP | HNRNPM | HepG2 | ENCSR979EWD | eCLIP | STAU2 | HepG2 | ENCSR006OEQ | eCLIP | FAM120A | K562 |
| ENCSR196INN | eCLIP | RBM15 | K562 | ENCSR586DGV | eCLIP | ZNF800 | K562 | ENCSR755TJC | eCLIP | HNRNPUL1 | HepG2 |
| ENCSR571ROL | eCLIP | XRCC6 | HepG2 | ENCSR485QCG | eCLIP | BCCIP | HepG2 | ENCSR773KRC | eCLIP | SRSF9 | HepG2 |
| ENCSR366YOG | eCLIP | QKI | K562 | ENCSR279UJF | eCLIP | SF3B4 | HepG2 | ENCSR724RDN | eCLIP | HNRNPL | HepG2 |
| ENCSR412NOW | eCLIP | HNRNPM | K562 | ENCSR589YHM | eCLIP | HLTF | K562 | ENCSR916SRV | eCLIP | TBRG4 | HepG2 |
| ENCSR135VMS | eCLIP | LSM11 | HepG2 | ENCSR893NWB | eCLIP | GRWD1 | HepG2 | ENCSR744GEU | eCLIP | IGF2BP1 | HepG2 |
| ENCSR661ICQ | eCLIP | PUM2 | K562 | ENCSR795CAI | eCLIP | HNRNPL | K562 | ENCSR550DVK | eCLIP | HNRNPC | HepG2 |
| ENCSR893EFU | eCLIP | DDX6 | K562 | ENCSR200DKE | eCLIP | MTPAP | K562 | ENCSR543TPH | eCLIP | AGGF1 | HepG2 |
| ENCSR059CWF | eCLIP | SBDS | K562 | ENCSR805SRN | eCLIP | LARP4 | HepG2 | ENCSR887FHF | eCLIP | FASTKD2 | K562 |
| ENCSR947JVR | eCLIP | DGCR8 | K562 | ENCSR845VGB | eCLIP | DDX55 | HepG2 | ENCSR819XBT | eCLIP | AATF | K562 |
| ENCSR356ZMO | eCLIP | AKAP1 | HepG2 | ENCSR023UHL | eCLIP | FASTKD2 | HepG2 | ENCSR506UPY | eCLIP | SUGP2 | HepG2 |
| ENCSR922WJV | eCLIP | PCBP1 | K562 | ENCSR265ZIS | eCLIP | GTF2F1 | HepG2 | ENCSR489ABS | eCLIP | RBM5 | HepG2 |
| ENCSR975KIR | eCLIP | IGF2BP1 | K562 | ENCSR486YGP | eCLIP | FUBP3 | HepG2 | ENCSR351PVI | eCLIP | SLTM | HepG2 |
| ENCSR177QQY | eCLIP | AKAP1 | K562 | ENCSR290VLT | eCLIP | MATR3 | HepG2 | ENCSR887LPK | eCLIP | EWSR1 | K562 |
| ENCSR766FAC | eCLIP | RPS3 | HepG2 | ENCSR993FMY | eCLIP | TROVE2 | HepG2 | ENCSR820DQJ | eCLIP | NOL12 | HepG2 |
| ENCSR539BEV | eCLIP | UPF1 | HepG2 | ENCSR295OKT | eCLIP | RBM22 | K562 | ENCSR981WKN | eCLIP | PTBP1 | K562 |
| ENCSR046JHH | eCLIP | CPEB4 | K562 | ENCSR769UEW | eCLIP | HNRNPA1 | HepG2 | ENCSR532VUB | eCLIP | CPSF6 | K562 |
| ENCSR023PKW | eCLIP | EIF3G | K562 | ENCSR154HRN | eCLIP | HNRNPA1 | K562 | ENCSR061SZV | eCLIP | DGCR8 | HepG2 |
| ENCSR121GQH | eCLIP | SERBP1 | K562 | ENCSR830BSQ | eCLIP | BUD13 | HepG2 | ENCSR384KAN | eCLIP | PTBP1 | HepG2 |
| ENCSR366DGX | eCLIP | KHSRP | HepG2 | ENCSR456JJQ | eCLIP | RBM22 | HepG2 | ENCSR958FKZ | eCLIP | PABPC4 | K562 |
| ENCSR141OIM | eCLIP | DDX6 | HepG2 | ENCSR238CLX | eCLIP | GEMIN5 | K562 | ENCSR739VVT | eCLIP | APOBEC3C | K562 |
| ENCSR520BZQ | eCLIP | HNRNPU | K562 | ENCSR438KWZ | eCLIP | ILF3 | K562 | ENCSR038JME | eCLIP | WRN | K562 |
| ENCSR432XUP | eCLIP | SRSF1 | K562 | ENCSR488JKQ | eCLIP | UTP18 | HepG2 | ENCSR921SXC | eCLIP | XPO5 | HepG2 |
| ENCSR534YOI | eCLIP | PRPF8 | K562 | ENCSR018WPY | eCLIP | AQR | HepG2 | ENCSR085JPB | eCLIP | WDR43 | HepG2 |
| ENCSR529GSJ | eCLIP | DHX30 | K562 | ENCSR964VOX | eCLIP | UTP18 | K562 | ENCSR734ZHL | eCLIP | UTP3 | K562 |
| ENCSR373ODC | eCLIP | SMNDC1 | HepG2 | ENCSR655NZA | eCLIP | XRN2 | HepG2 | ENCSR815VVI | eCLIP | CDC40 | HepG2 |
| ENCSR867DSZ | eCLIP | NPM1 | K562 | ENCSR725ARB | eCLIP | AGGF1 | K562 | ENCSR061EVO | eCLIP | SND1 | HepG2 |
| ENCSR834YLD | eCLIP | DROSHA | HepG2 | ENCSR668MJX | eCLIP | GRSF1 | HepG2 | ENCSR121NVA | eCLIP | PRPF8 | HepG2 |
| ENCSR268ETU | eCLIP | HNRNPK | K562 | ENCSR570WLM | eCLIP | QKI | HepG2 | ENCSR844RVX | eCLIP | EFTUD2 | K562 |
| ENCSR081JYH | eCLIP | NSUN2 | K562 | ENCSR018ZUE | eCLIP | FKBP4 | HepG2 | ENCSR506OTC | eCLIP | TBRG4 | K562 |
| ENCSR339FUY | eCLIP | PCBP2 | HepG2 | ENCSR269AJF | eCLIP | RPS11 | K562 | ENCSR197INS | eCLIP | PPIL4 | K562 |
| ENCSR923NKN | eCLIP | DDX55 | K562 | ENCSR440SUX | eCLIP | MATR3 | K562 | ENCSR861PAR | eCLIP | NONO | K562 |
| ENCSR970NKP | eCLIP | LIN28B | K562 | ENCSR903PRV | eCLIP | FTO | HepG2 | ENCSR987FTF | eCLIP | RBFOX2 | HepG2 |
| ENCSR301UQM | eCLIP | GNL3 | K562 | ENCSR256CHX | eCLIP | PCBP1 | HepG2 | ENCSR224QWC | eCLIP | FXR2 | K562 |
| ENCSR484LTQ | eCLIP | NCBP2 | K562 | ENCSR961OKA | eCLIP | LARP7 | HepG2 | ENCSR663WES | eCLIP | BUD13 | K562 |
| ENCSR888YTT | eCLIP | LARP4 | K562 | ENCSR565DGW | eCLIP | DHX30 | HepG2 | ENCSR303OQD | eCLIP | METAP2 | K562 |
| ENCSR018RVZ | eCLIP | NCBP2 | HepG2 | ENCSR120EAR | eCLIP | RPS3 | K562 | ENCSR206RXT | eCLIP | AKAP8L | K562 |
| ENCSR097NEE | eCLIP | PPIG | HepG2 | ENCSR663NRA | eCLIP | ZRANB2 | HepG2 | ENCSR893RAV | eCLIP | U2AF2 | K562 |
| ENCSR930BZL | eCLIP | DDX3X | K562 | ENCSR786TSC | eCLIP | ILF3 | HepG2 | ENCSR041NUV | eCLIP | EIF3D | HepG2 |
| ENCSR965DLL | eCLIP | SFPQ | HepG2 | ENCSR001VAC | eCLIP | NOLC1 | K562 | ENCSR657TZB | eCLIP | XRN2 | K562 |
| ENCSR277DEO | eCLIP | NKRF | HepG2 | ENCSR308YNT | eCLIP | PUM1 | K562 | ENCSR916XIV | eCLIP | EIF3H | HepG2 |
| ENCSR202BFN | eCLIP | U2AF2 | HepG2 | ENCSR128VXC | eCLIP | SND1 | K562 | ENCSR736AAG | eCLIP | GTF2F1 | K562 |
| ENCSR202HKN | eCLIP | WDR3 | K562 | ENCSR647HOX | eCLIP | HLTF | HepG2 | ENCSR647CLF | eCLIP | GPKOW | K562 |
| ENCSR774RFN | eCLIP | FXR1 | K562 | ENCSR606BPV | eCLIP | AQR | K562 | ENCSR301TFY | eCLIP | DKC1 | HepG2 |
| ENCSR658IQB | eCLIP | SMNDC1 | K562 | ENCSR876EYA | eCLIP | BCLAF1 | HepG2 | ENCSR214BZA | eCLIP | DDX59 | HepG2 |
| ENCSR349CMI | eCLIP | WDR43 | K562 | ENCSR438GZQ | eCLIP | KHSRP | K562 | ENCSR820WHR | eCLIP | POLR2G | HepG2 |
| ENCSR040QLV | eCLIP | DDX21 | K562 | ENCSR756CKJ | eCLIP | RBFOX2 | K562 | ENCSR490IEE | eCLIP | UCHL5 | HepG2 |
| ENCSR891RIC | eCLIP | NIPBL | K562 | ENCSR693JWP | eCLIP | EXOSC5 | HepG2 | ENCSR291XPT | eCLIP | PUS1 | K562 |
| ENCSR356MSW | eCLIP | SSB | K562 | ENCSR307YIW | eCLIP | EIF4G2 | K562 | ENCSR943MHU | eCLIP | SAFB2 | K562 |
| ENCSR735HOK | eCLIP | YBX3 | HepG2 | ENCSR349KMG | eCLIP | UCHL5 | K562 | ENCSR314UMJ | eCLIP | TRA2A | HepG2 |
| ENCSR571VHI | eCLIP | HNRNPUL1 | K562 | ENCSR712IAG | eCLIP | ZC3H11A | K562 | ENCSR328LLU | eCLIP | U2AF1L5,U2AF1 | HepG2 |
| ENCSR580MFX | eCLIP | SUPV3L1 | HepG2 | ENCSR361OCV | eCLIP | NIP7 | HepG2 | ENCSR989SMC | eCLIP | FTO | K562 |
| ENCSR527DXF | eCLIP | EFTUD2 | HepG2 | ENCSR623VEQ | eCLIP | TIA1 | HepG2 | ENCSR907GUB | eCLIP | ZC3H11A | HepG2 |
| ENCSR468FSW | eCLIP | SRSF7 | K562 | ENCSR069EVH | eCLIP | FUS | K562 | ENCSR628IDK | eCLIP | KHDRBS1 | K562 |
| ENCSR483NOP | eCLIP | SLBP | K562 | ENCSR057DWB | eCLIP | TIA1 | K562 | ENCSR539ZTS | eCLIP | TROVE2 | K562 |
| ENCSR000SSH | eCLIP | SLTM | K562 | ENCSR580OFI | eCLIP | SUPV3L1 | K562 | ENCSR133QEA | eCLIP | SF3B1 | K562 |
| ENCSR919HSE | eCLIP | CSTF2T | HepG2 | ENCSR657TZZ | eCLIP | ZNF622 | K562 | ENCSR013CTQ | eCLIP | EXOSC5 | K562 |
| ENCSR576SHT | eCLIP | DDX42 | K562 | ENCSR568DZW | eCLIP | TAF15 | K562 | ENCSR456ASB | eCLIP | UPF1 | K562 |
| ENCSR513NDD | eCLIP | SRSF7 | HepG2 | ENCSR258QKO | eCLIP | XRCC6 | K562 | | | | |
| ENCSR825SVO | eCLIP | AARS | K562 | ENCSR484LAB | eCLIP | SAFB | K562 | | | | |
